# Cholesterol deprivation induces TGFβ signaling to promote basal differentiation in pancreatic cancer

**DOI:** 10.1101/633719

**Authors:** Linara G. Cornell, Suraj Peri, Diana Restifo, Alena Klochkova, Tiffiney R. Hartman, Alana M. O’Reilly, Ralph Francescone, Janusz Franco-Barraza, Neelima Shah, Emmanuelle Nicolas, Elizabeth A. Handorf, Kathy Q. Cai, Alexandra Mazitova, Ido Sloma, Rachel Chiaverelli, Richard Moffitt, Erica A. Golemis, Edna Cukierman, Igor Astsaturov

**Author notes:** Contact information: Igor Astsaturov, Fox Chase Cancer Center, 333 Cottman Avenue, Philadelphia, PA 19111.

## Abstract

Oncogenic transformation alters the metabolism of cellular nutrients to sustain tumor growth. We here define a mechanism by which modifications in cholesterol metabolism control the formation of pancreatic ductal adenocarcinoma (PDAC). Disruption of distal cholesterol biosynthesis by means of conditional inactivation of *Nsdhl* in mice bearing a tumor-inducing *Kras* mutation (*Kras^G12D^*) prevented PDAC formation in the context of a heterozygous *Trp53^f/+^*genotype without impairing normal pancreatic development. In mice with pancreatic *Nsdhl* ablation and homozygous loss of *Trp53*, the emerging tumors presented with the aggressive basal (mesenchymal) phenotype as opposed to the classic (glandular) PDAC. This paralleled significantly reduced expression of cholesterol metabolic pathway genes in human basal PDAC subtype. Mechanistically, we demonstrate that genetic or metabolic cholesterol deprivation stabilizes the transforming growth factor beta (TGFβ) receptor to activate pro-mesenchymal effectors in human and murine PDAC, providing a direct mechanism by which cholesterol metabolism can condition tumor differentiation.

## Introduction

The incidence of pancreatic ductal adenocarcinoma (PDAC) has been rising for the past decades, and by 2030, it is poised to become the second leading cause of cancer deaths in the USA (Chari et al., 2015; Rahib et al., 2014). The nearly identical numbers of PDAC diagnoses and PDAC deaths have been associated with high metastatic propensity of this cancer. Epithelial-to-mesenchymal transition (EMT) has been proposed as one of the key mechanisms of drug resistance (Sabnis and Bivona, 2019; Zheng et al., 2015), cancer metastatic dissemination and invasive growth (Aiello et al., 2017).

Gene expression analyses have established at least two molecular subtypes of PDAC, the classic (also known as glandular), and the basal (i.e., mesenchymal (Aung et al., 2018; Moffitt et al., 2015)), each of which is associated with distinct prognoses and sensitivity to chemotherapy, so that the median survival of basal PDAC is nearly half of that in classic subtype (Aung et al., 2018). The defining features of the basal subset of PDAC include increased signaling by transforming growth factor beta (TGFβ), and expression of TGFβ-induced genes associated with a mesenchymal phenotype, such as transcription factors ZEB1, ZEB2, TWIST, SNAI1, and SNAI2 (Scheel et al., 2011), as well as reduced expression of epithelial adhesion molecule E-cadherin (CDH1) and epithelial lineage transcription factors GATA6, SOX17, and HNF4A (Bailey et al., 2016; Collisson et al., 2011; Moffitt et al., 2015; TCGA., 2017). Although TGFβ signaling plays a central role in EMT of established tumors, it has a potent tumor suppressive role in untransformed cells, as reflected by frequent inactivation of components of the TGFβ pathway (TGFBR1, TGFBR2, SMAD4) in about a quarter of PDAC cases (David et al., 2016; Stankic et al., 2013). At present, little is known of the epigenetic or metabolic factors that act upstream of TGFβ in EMT, and influence PDAC evolution into the basal versus classic subtypes.

In epidemiological studies, the rising incidence of PDAC has been linked to elevated serum cholesterol (dyslipidemia) and obesity (Genkinger et al., 2015), which are attributable in part to high fat and calorie Western diets. Oncogene-transformed cancer cells accelerate uptake and endogenous biosynthesis of cholesterol and phospholipids (Pitroda et al., 2009; Silvente-Poirot and Poirot, 2014). In mouse models of PDAC, driver mutations in the oncogene *Kras* coupled with loss of the tumor suppressor *Trp53* reprogram cellular metabolism in numerous ways affecting utilization of energy sources, accelerating cholesterol biosynthesis and uptake (Freed-Pastor et al., 2012; Ying et al., 2012). Although cholesterol has been proposed to drive PDAC carcinogenesis, this idea has been challenged by the relatively modest clinical activity of cholesterol-lowering medications such as statins aimed to reduce PDAC risk (Bang et al., 2018). Instead, the lack of correlation between PDAC mortality and serum cholesterol levels (Huang et al., 2017) raises the possibility of a tumor cell-centric mechanism by which cholesterol can modulate oncogenic functions and modify disease course at earlier stages of PDAC formation. Supporting this idea, data implicating cholesterol and its metabolites in cancer initiation and progression are beginning to emerge (Gabitova et al., 2015; Nelson et al., 2013). In one notable example, liver X-receptor pathway activation, an important nodule of the cholesterol metabolism, in a mouse model of breast cancer in conjunction with genetically induced accumulation of 27-hydroxycholesterol, fostered the development of aggressive basal adenocarcinoma with mesenchymal differentiation. These findings suggested that the network of genes and metabolites regulating cholesterol homeostasis can directly condition cancer differentiation (Nelson et al., 2013). Despite these advances, the mechanism for this dramatic effect of cholesterol metabolism on cancer epigenetic program remains to be defined.

In this work, we employed a genetic mouse model to explore cholesterol dependency in PDAC. The enzymatic activity of NSDHL (NAD(P)-dependent steroid dehydrogenase-like) mediates an irreversible step in the oxidative decarboxylation of two methyl moieties at the C4 position of a cholesterol precursor, which is essential for endogenous cholesterol biosynthesis (Cunningham et al., 2015; Gabitova et al., 2015). Conditional inactivation of *Nsdhl* in pancreatic epithelial cells via *Pdx1-Cre* transgene enabled us to study the role of endogenous cholesterol biosynthesis in the context of pancreatic cancer model combining *<u>C</u>re*-activated oncogenic *<u>K</u>ras^G12D^* allele and conditional knockout of “floxed” *Tr<u>p</u>53* (aka, *KPC* mice, (Bardeesy et al., 2006)). Remarkably, genetically induced cholesterol auxotrophy in the pancreatic epithelium blocked malignant progression of pancreatic precursor lesions and protected *Trp53^+/−^*mice from PDAC development. In unexpected contrast, in the context of bi-allelic rather than heterozygous inactivation of *Trp53*, loss of *Nsdhl* promoted basal phenotype of *Kras^G12D^*-induced PDAC. This metabolically-determined dichotomy of PDAC differentiation was mediated by cholesterol-sensitive regulation of TGFβ signaling. Importantly, the results that we obtained from animal model paralleled in our analyses the human PDAC transcriptome data in which cholesterol pathway mRNA signatures are strikingly different and independently prognostic when comparing between classic and basal subtypes. Finally, we define a mechanism by which cholesterol abundance controls TGFβ signaling. Contingent on complete p53 inactivation and under cancer cell cholesterol auxotrophy, cellular cholesterol regulated TGFβ ligand abundance and dynamics of receptor activation via canonical SMAD2/3 effector cascade, promoted the EMT transcriptional program to attain an aggressive and poorly differentiated, basal, PDAC phenotype.

## Results

### Disruption of epithelial cholesterol biosynthesis prevents PDAC development

To test dependency of pancreatic carcinogenesis on tumor-intrinsic cholesterol biosynthesis, we used mice in which conditional inactivation of *Nsdhl* via a *Pdx1*-*Cre* transgene is expected to take place in the developing pancreatic bud at embryonic day 8.5 (Gabitova et al., 2015; Offield et al., 1996). The efficiency of *Nsdhl* inactivation in pancreatic epithelium was evident from complete Cre-mediated *Nsdhl^f/f^*genomic rearrangement and loss of detectable NSDHL expression in pancreatic tissue (*Nsdhl*^*ΔPanc*^ mice, **Fig. S1A, B**). These *Nsdhl*^*ΔPanc*^ mice produced normal-sized litters at term with an approximate 1:1 male to female ratio; given the location of *Nsdhl* on the X chromosome, this implied no embryonic lethality. Comparison of pancreatic tissues of *Nsdhl*^*ΔPanc*^ and wild type mice at 6-8 weeks of age demonstrated normal pancreas size (**Fig. S1C**), normal pancreatic ductal and acinar anatomy (**Fig. S1D**), and proliferation rate (**Fig. S1E**). To probe the regenerative capacity of the *Nsdhl*-deficient pancreas, we treated adult *Nsdhl*^*ΔPanc*^ and wild type animals with a two-day course of caerulein, an established inducer of acute pancreatitis (Renner et al., 1983). We saw no significant morphological differences during acinar regeneration following acute caerulein-induced injury, and no differences in proliferation (Ki67, **Fig. S1F, G**), suggesting normal regenerative capacity of *Nsdhl*^*ΔPanc*^ epithelial cells.

We next evaluated the effects of conditional *Nsdhl* knockout in an established model of pancreatic carcinogenesis induced by oncogenic *Kras^G12D^* (*KC* mice) in which mice develop pre-malignant lesions by 6 months of age (Hingorani et al., 2005; Jackson et al., 2001; Tuveson et al., 2004). *KC* and *KCN* animals (denoting the *LSL-<u>K</u>ras^G12D^;Pdx1*-*<u>C</u>re* genotype with or without *<u>N</u>sdhl^f/f^* allele, respectively) developed pre-malignant epithelial lesions represented by acinar-to-ductal metaplasia (ADM) and low grade pancreatic intraepithelial neoplasia (PanIN) at a similar low preponderance per pancreas (**Fig. S2A**). These results indicate that arresting endogenous epithelial cholesterol biosynthesis, at the level of NSDHL, is insufficient to counter the increased proliferative effect of *Kras^G12D^*seen in pancreatic epithelial cells.

Progression of PanIN to PDAC, in the pancreas of both mice and humans, requires inactivation of the P53 tumor suppressor in conjunction with constitutive KRAS activity (Hingorani et al., 2005). Control animals with monoallelic loss of *Trp53* (denoted as *Pdx1-Cre;LSL-Kras^G12D^;Trp53^f/+^*, or *KPC* mice) developed pancreatic adenocarcinoma starting from 2 months of age with a medium overall survival of about 5 months (**Fig. 1A**). The majority of evaluable PDAC in these *KPC* animals were represented by grade 1 and 2 adenocarcinoma with glandular differentiation features (**Table S1**). In contrast, the *KPCN* mice developed PDAC at much later term and only in 3 out of 37 animals (**Fig. 1A**). This translated to a markedly superior PDAC-free survival at 6 months of age in *KPCN* compared to *KPC* (92% vs. 30%, respectively, log rank test *p*<0.0001), despite a similar preponderance of proliferating Ki67^+^ cells (**Fig. S2B**).

**Figure 1.**
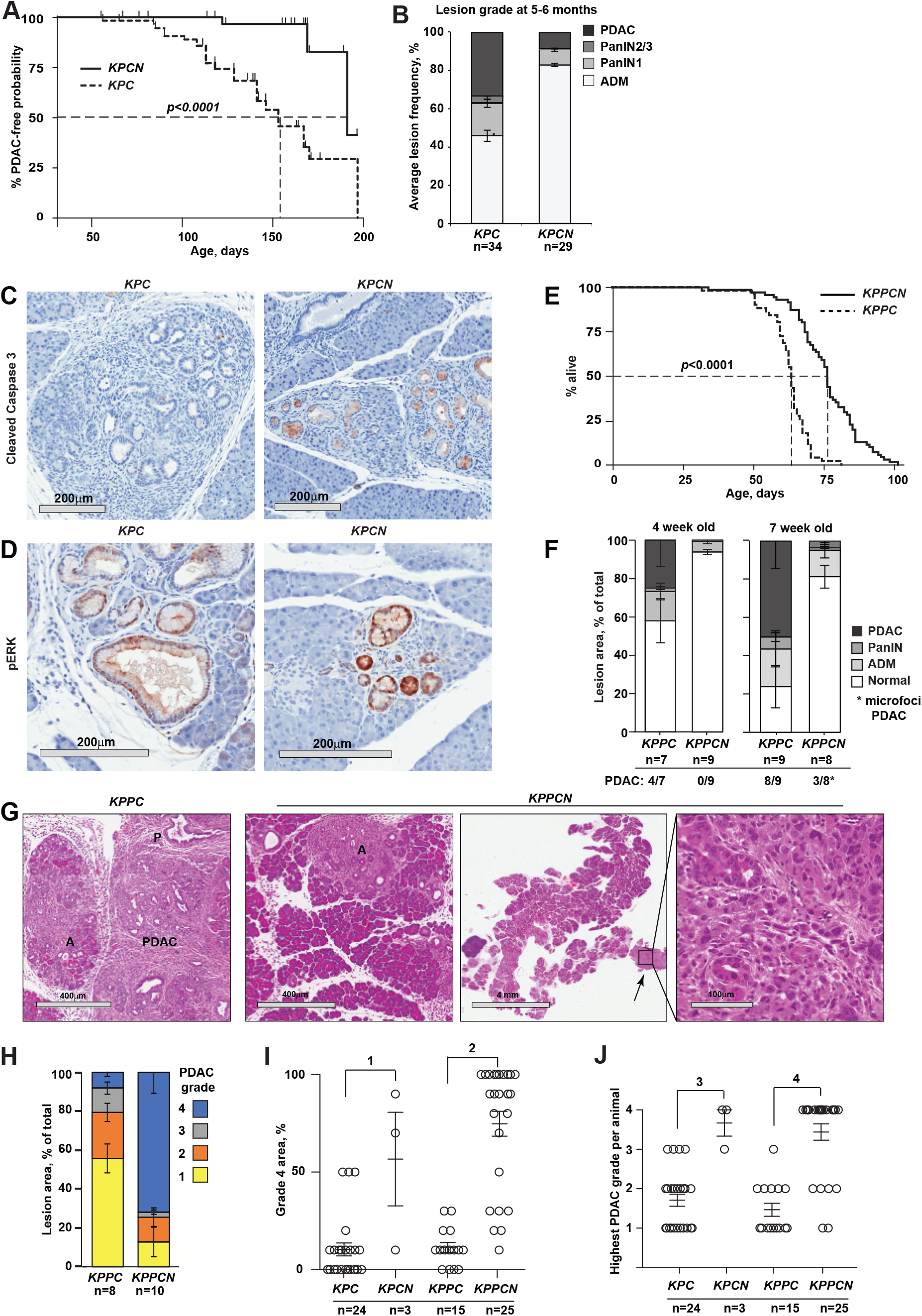
Effects of NSDHL deficiency on pancreatic adenocarcinoma development. (**A**) Kaplan-Meier PDAC-free survival of *KPC* (n=34) and *KPCN* (n=37) mice. *p*<0.0001, Logrank test. (**B**) Frequency of pancreatic epithelial lesion by grade per section in *KPC* and *KPCN* mice at 5-6 months of age; *p*=0.035 for PDAC, Fisher’s exact test; *p*=0.02 for PanIN2/3; *p*=0.0006 for ADM, Wilcoxon test. (**C, D**) Expression of cleaved caspase 3 (**C**) and phosphorylated ERK1/2 (**D**) in pancreatic ADM and PanIN lesions. (**E**) Kaplan-Meier survival of *KPPC* (n=64) and *KPPCN* (n=76) mice. *p*<0.0001, Logrank test. (**F**) Delayed progression of pancreatic epithelial lesion in *KPPCN* mice compared to age-matched *KPPC* at 4 and 7 weeks shown as relative lesion area per section; *p*<0.02 for comparisons of PDAC and normal areas, Wilcoxon test. (**G**) Hematoxylin and eosin stained pancreas sections at 7 weeks of age of *KPPC* and *KPPCN* mice, respectively. Small foci of grade 4 PDAC (*arrow*) are seen on the background of nearly normal *KPPCN* pancreatic tissue with ADM (A) and early PanIN (P) lesions. Scale bars, as shown. (**H**) Histological grading of advanced PDAC tumors; *p<*0.01 for grades 1, 3 and 4. (**I**) Comparison of grade 4 PDAC areas per section; (1) and (2), *p*<0.0001. (**J**) Comparisons of the highest PDAC grades per animal; (3) *p=*0.0009. In all figures, data are represented as mean±SEM, *p*-values by Wilcoxon test. See also Figure S2.

Histological evaluation of pancreatic tissues of age-matched mice at 5-6 months demonstrated a correspondingly low burden of advanced PanIN and PDAC lesions in *KPCN* mice (**Fig. 1B**) compared to *KPC*. Further, *KPCN* pancreatic tissues exhibited a robust expression of cleaved caspase 3 within ADM and PanIN lesions, compared to *KPC* (**Fig. 1C, S2C**), which were not observed in normally appearing acinar cells (**Fig. 1C**). We further investigated possible effects of NSDHL inactivation on the signaling downstream of KRAS and found slightly increased expression of phosphorylated form of MAPK1/2 in *KPCN* pancreatic premalignant lesions (**Fig. 1D, S2D**) indicating no inhibitory effects of NSDHL deficiency on proliferation-associated MAPK signaling. The phosphorylated ribosomal protein S6, an mTOR kinase target and a correlate of metabolic activity (Peterson and Sabatini, 2005), tended to be diminished in *KPCN* PanIN lesions (**Fig. S2E**).

Deficiency of *Nsdhl* has been reported to cause accumulation of C4-methylsterols, a group of metabolites arising from cholesterol pathway inhibition that activate the transcription factor LXRα (Gabitova et al., 2015). Confirming significant inhibition of cholesterol biosynthesis in the *KPCN* pancreas, we found significantly elevated expression of the LXRα–dependent transcript ATP-binding cassette transporter member 1 (ABCA1) in ADM and PanIN lesions, in contrast to *KPC* (**Fig. S2F**). Notably, detailed examination demonstrated that the small number of PDAC arising in KPCN mice were of poorly differentiated histology (**Table S1**).

### Cholesterol restriction promotes loss of differentiation and epithelial-mesenchymal transition (EMT) in Kras-induced PDAC in the context of complete Trp53 loss

In the *KPPC* model (*Pdx1-Cre;LSL-Kras^G12D^;Trp53^f/f^*), complete loss of *Trp53* induces rapid development of PDAC (Bardeesy et al., 2006). We next compared mice with bi-allelic inactivation of *Trp53*, with or without conditional knockout of *Nsdhl^f/f^* (designated *KPPC* or *KPPCN*) (**Fig. 1E**). Inactivation of NSDHL in the absence of P53 only marginally improved the median survival of *KPPCN* compared to approximately 2 months attained in *KPPC* despite the markedly delayed progression of premalignant lesions to PDAC in *KPCN* mice (**Fig. 1A, B**). Most *KPPCN* animals commenced significant weight loss around day 70, resulting in a relatively small although statistically significant difference in median survival compared to *KPPC* mice (**Fig. 1E,** median survival, 76 vs. 64 days, respectively; *p*<0.0001, log rank test).

To investigate the pathological features of *KPPCN* carcinomas, we evaluated pancreatic tissues at 4 and 7 weeks of age which surprisingly showed a markedly reduced burden of advanced PanIN pancreatic lesions and PDAC in the *KPPCN* compared to *KPPC* group (**Fig. 1F**). At 4 weeks, PDAC was detectable in 4/7 *KPPC* pancreatic tissues, while no evidence of tumors was apparent in the age-matched *KPPCN* animals. At 7 weeks, 8 of 9 *KPPC* mice had developed multifocal large pancreatic tumors comprising more than 50% of the organ area. At the same age, only 3 of 8 *KPPCN* mice had developed microscopic PDAC foci, comprising <5% of the pancreas (**Fig. 1F**). Importantly, these microscopic foci were comprised of poorly differentiated adenocarcinoma (indicated by arrow, **Fig. 1G**) and appeared on the background of low grade PanIN and ADM lesions. Micro-magnetic resonance imaging (mMRI) of larger cohorts demonstrated advanced large PDAC tumors and a typically enlarged pancreas in 7 week old *KPPC* mice, while *KPPCN* mice typically had a normally sized pancreas (**Fig. S3A**), which was also reflected in a marked difference in pancreas weight between the two genotypes (**Fig. S3B**). We next compared pathological features of advanced PDAC tumors present in *KPPC* and *KPPCN* mice at the time of euthanasia. Whereas in *KPPC* mice PDAC were typically grade 1-2, tumors in *KPPCN* mice demonstrated nearly uniform development of grade 4 PDAC (**Fig. 1H-J, Fig. S3C** for grading, and **Table S1**), echoing the decreased differentiation of *KPCN* versus *KPC* tumors (**Table S1**).

Analyzing tumor differentiation in greater depth, *KPPC* tumors were characterized by cytokeratin-positive pseudoglandular structures with strong expression of E-cadherin (CDH1 in **Fig 2A**), as was previously established for this model of PDAC (Bardeesy et al., 2006). In contrast, the majority of *KPPCN* carcinomas were comprised of single cells or small clusters of spindle-shaped tumor cells with weak expression of cytokeratin and absence of CDH1 expression (**Fig 2B**). The *KPPCN* carcinomas also had a lower density of CD31+ blood vessels (**Fig. S3D, E**), ruling out an *Nsdhl*-dependent angiogenesis switch previously implicated in PDAC progression (Rhim et al., 2014). Since loss of CDH1 expression is a feature of EMT (Scheel et al., 2011)), we further examined the differentiation markers of *KPPCN* carcinomas. Direct labeling of cells isolated from tumor tissues with EPCAM and PDGFRα antibodies allowed us to discern the populations of epithelial carcinoma cells and cancer-associated fibroblasts (CAFs) (**Fig. S4A**). Fluorescence-activated cell sorting (FACS) of cells isolated from the spontaneous pancreatic tumors showed significant reduction in EPCAM-positive cells in *KPPCN* as compared to *KPPC* (10% vs. 50%, respectively; **Fig. 2C, D**). This reduction could either reflect the possibility that *KPPCN* tumors were disproportionally composed of CAFs, or alternatively could indicate that a significant number of *KPPCN* cancer cells had lost epithelial lineage markers. The cultures of FACS-isolated cellular populations from *KPPCN* tumors (predominantly PDGFRα-/EPCAM-double-negative, or PDGFRα+) carried Cre-mediated rearrangements of *Kras* and *Nsdhl* indicating epithelial origin of these cells (**Fig. S4B-E**).

**Figure 2.**
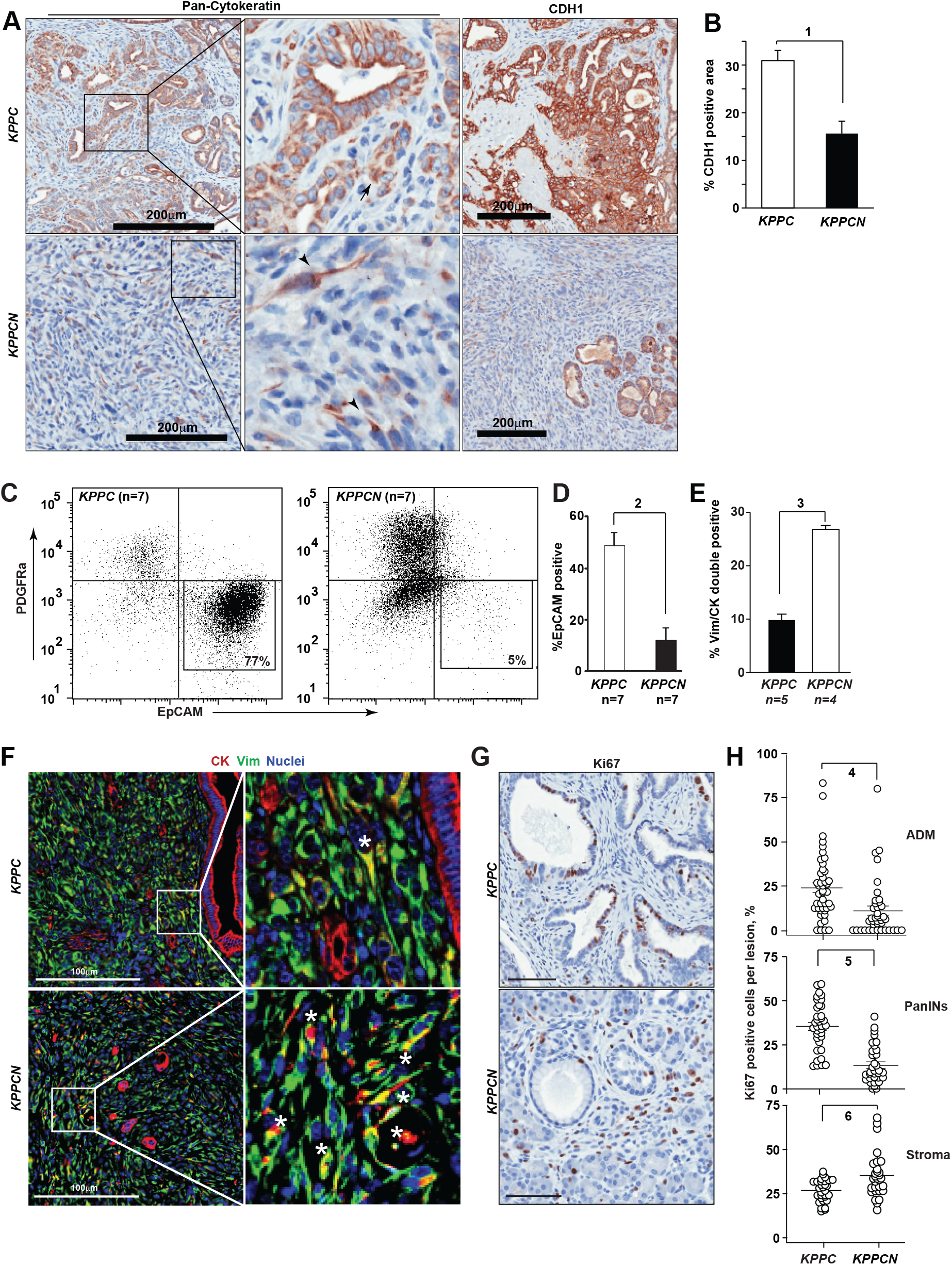
Development of basal pancreatic adenocarcinoma with mesenchymal differentiation in *Nsdhl* knockouts. (**A**) Representative pan-cytokeratin (CK) and E-cadherin (CDH1) immunohistochemical staining of pancreatic tumor tissues. *Scale bars*, 200 µm. (**B**) Quantification of CDH1 positive areas; (1) *p*=0.001. (**C**) Fluorescent activated cell sorting (FACS) of primary pancreatic tumor tissue shows markedly lower percentage of EPCAM+ cells in *KPPCN* compared to *KPPC*. CD45+ myeloid cells were excluded by gating. (**D**) Percentage of EPCAM-positive epithelial cells assessed by FACS in primary *KPPC* and *KPPCN* tumors; (2) *p*=0.0012, Wilcoxon test. (**E**) Quantification of CK+/VIM+ double positive cells by multicolor immunofluorescence; (2) *p*=0.016, Wilcoxon test. (**F**) Representative images of multiplex immunofluorescence characterization of *KPPC* and *KPPCN* carcinoma, depicting cytokeratin (CK) positive cells in red, vimentin (Vim) positive cells in green, and nuclei in blue. *Asterisks*, denote double-positive (CK+, VIM+) cells (**G**) Quantification of Ki67+ cells in pancreatic ADM, PanIN lesions and surrounding stroma in *KPPC* (n=7) and *KPPCN* (n=9) 4-week old mice. Data are represented as percent of Ki67+ in individual lesions; (4) *p*=0.001; (5) *p*<0.0001; (6) *p*=0.0015, Student t-test. (**H**) Representative Ki67 expression in stroma and epithelial lesions. In all figures, data are represented as mean±SEM. See also Figures S2-S4.

In keeping with the established pathology for the murine pancreatic adenocarcinoma induced by oncogenic *Kras^G12D^*and *Trp53* inactivation (Aiello et al., 2018; Bardeesy et al., 2006; Hingorani et al., 2005), the *KPPC* tumors exhibited glandular CK+ and CDH1+ epithelial structures (**Fig. 2A, F**) clearly separated from the cancer-associated fibroblasts expressing vimentin (VIM). As an alternative approach to assessing the interaction of the epithelial and the stromal compartmentalization of *KPPC* and *KPPCN* tumors, we used immunofluorescent labeling of cytokeratin (CK) and VIM markers, respectively. Only a small fraction (<10%, **Fig. 2E**) of cells in *KPPC* tumor co-expressed VIM and CK; these were typically single cells or small clusters of carcinoma cells localized outside of larger glandular structures (**Fig. 2F**). In contrast, in *KPPCN* tumors the overall frequency of double-positive VIM+/CK+ cells was significantly elevated (**Fig. 2F**), with these cells comprising the bulk of the tumor. We conclude that the higher prevalence of VIM+/CK+ double positive cells and loss of CDH1 expression in *KPPCN* tumors reflects their mesenchymal differentiation, consistent with the reported basal subtype of PDAC(Moffitt et al., 2015).

Independent of the *KPPCN* tumors adapting a more mesenchymal morphology, these tumors might differentially condition the essential tumor-supporting activity of pancreatic CAFs. The heterogeneous CAF population in pancreatic cancer is comprised of secretory fibroblasts and α-smooth muscle actin-positive (αSMA+) CAFs (also known as myofibroblasts) (Ohlund et al., 2017). The latter are responsible for enhanced deposition of collagen I, a characteristic of fibrotic desmoplasia in classic PDAC, and have been suggested to restrain PDAC progression in some models(Ozdemir et al., 2014). Assessment of the stroma of *KPPCN* tumors showed markedly reduced collagen deposition and fibrosis, based on use of type I collagen-directed antibodies and by Masson’s trichrome staining, respectively (**Fig. S5A, B**), and paucity of αSMA+ myofibroblasts (**Fig. S5C, D**), altogether indicating that basal PDAC induced an expansion of CAF populations expressing high level of GLI1 (**Fig. S5E, F**) and phosphorylated focal adhesion kinase (pY397-FAK, **Fig. S5G, H**) compared to classic (i.e. glandular) PDAC(Ohlund et al., 2017). The activated state of the tumor-associated stroma was also apparent at the premalignant stage, as assessed by Ki-67 positivity in pancreatic stromal elements surrounding *KPPCN* ADM and PanIN lesions compared to *KPPC* (**Fig. 2G, H**). These results together suggest that development of basal PDAC is distinctly different from the canonical ADM-PanIN-PDAC linear progression in classic glandular subtype of pancreatic cancer, and is associated with hyperactivated stroma.

### KPPCN PDAC cell-intrinsic activation of TGFβ and other pro-EMT signaling

To understand cell-intrinsic differences in EMT-related signaling mechanisms distinguishing *KPPC* and KPPCN *PDAC*, we generated an extended panel of cell lines from *KPPC* (n=10) and *KPPCN* (n=10) tumors using differential trypsinization followed by FACS sorting enrichment of PDAC cells, which were analyzed at early passages (passage 3-5). The epithelial identity of these cell lines was validated by expression of CK, albeit sometimes low, and by completeness of *Kras* and *Trp53* gene rearrangements (**Fig. S6A, B**). *In vitro*, we observed about half of *KPPCN* cell lines grew in a pattern of scattered single cells suggesting an EMT phenotype (Scheel et al., 2011) (**Fig. S4C**). Importantly, the observed *in vitro* EMT phenotypes of *KPPCN* cells were stable and correlated with development of diffuse undifferentiated carcinomas following syngeneic orthotopic implantation and resulted in shorter survival of animals compared to similar implantations of *KPPC* PDAC cells (**Fig. S6C, D**).

Five of 10 KPCN and 1/10 KPC cell lines were classified as mesenchymal due to loss of CDH1 on the cell surface, as determined by flow cytometry (**Fig. S6E**). To identify the mechanistic basis of the enhanced EMT of *KPPCN* cancer cells, we analyzed mRNA from 3 independent PDAC clones derived from the *KPPC* and *KPPCN* tumors using next generation sequencing, focusing initially on lines with the typical phenotypes: epithelial for *KPPC* and mesenchymal for *KPPCN* (**Fig. 3A** and **Table S2**). Confirming a sustained cell-intrinsic effect of *Nsdh*l deficiency of lipid metabolism, *KPPCN* carcinoma cells showed globally suppressed expression of multiple genes involved in biosynthesis of phospholipids and cholesterol (**Fig. 3A, B**). In addition to genetically induced *Nshdl* deficiency, expression of the distal cholesterol biosynthesis pathway gene *Ebp* and *Dhcr24,* along with other canonical SREBP targets were markedly reduced (**Fig. 3B**).

**Figure 3.**
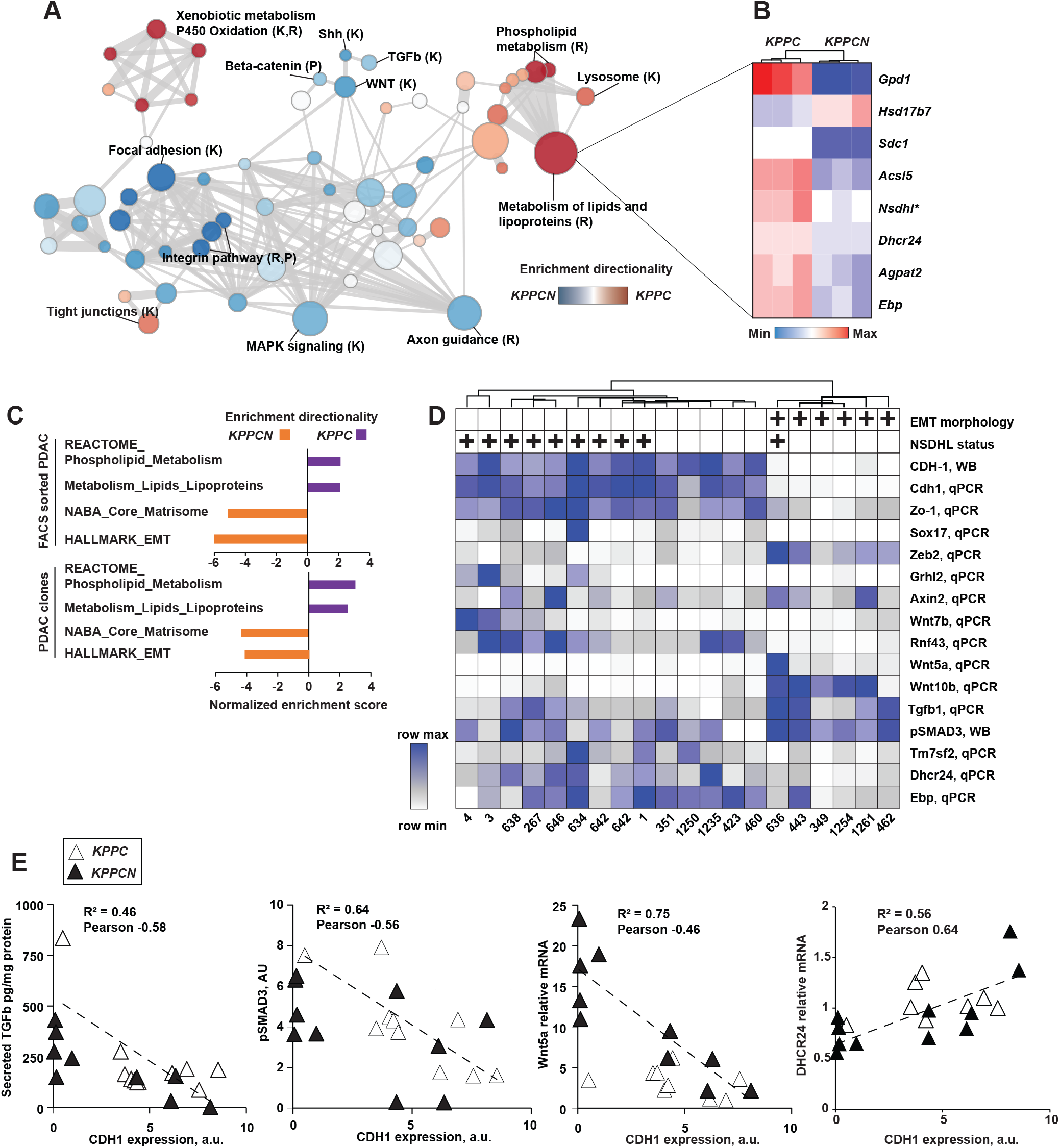
Conditional knockout of *Nsdhl* promotes epithelial-to-mesenchymal transition (EMT) switch in mouse pancreatic adenocarcinoma. (**A**) Gene Set Enrichment Analysis (GSEA) of differentially expressed genes as determined by mRNA sequencing of KPPC and KPPCN carcinoma clones, n=3 per genotype. Nodes sizes in the graph are proportionate to the number of genes in a given gene set, thickness of edges is proportionate to the number of shared genes between the gene sets. Sources of signatures: K, KEGG (Kyoto Encyclopedia of Genes and Genome); R, www.Reactome.org. (**B**) Heat map of a selected list of differentially expressed cholesterol and lipid metabolism genes. (**C**) Normalized enrichment scores for each genotype of FACS sorted primary PDAC cells and PDAC clones (shown, family-wise error rate <5%). (**D**) Unsupervised hierarchical clustering of quantified expression of indicated genes. qPCR, quantitative reverse transcription PCR; WB, Western blot. Mesenchymal morphology *in vitro* and NSDHL status is indicated above the heatmap by (*+*) positive. (**E**) Correlation of CDH1 expression and secreted TGFβ1, phosphorylated SMAD2/3, WNT5A and DHCR24 mRNA in *KPPC* and *KPPCN* clones. See also Figure S5 and Table S2.

Reciprocally, the *KPPCN* transcriptome was significantly enriched in gene targets associated with the EMT-promoting signaling pathways including TGFβ (Scheel et al., 2011), as well as PDGFR, integrin, and WNT, whereas expression of components of tight junctions (KEGG), a hallmark of epithelial differentiation, was enriched in the KPPC transcriptome (**Fig. 3A**). These differentially expressed mRNA signatures were similarly reproduced in RNA sequencing analyses of KPPC and KPPCN PDAC cells freshly isolated by FACS sorting from primary tumors (**Fig. 3C**); we observed highly significant overrepresentation of phospholipid and lipid metabolism gene transcripts in KPPC cells, and overrepresentation of signatures for EMT and extracellular matrix genes in KPPCN cells (**Fig. 3C** and **Table S2**).

We then used qRT-PCR to explore patterns of gene expression for key components of these pathways across the entire cell line panel (**Fig. 3D**). Among the *KPPCN* cell lines, those characterized as epithelial based on retention of *CDH1* clustered with 9/10 KPPC cell lines, implying loss of *Nsdhl* predominantly acted by increasing the likelihood of mesenchymal transition. Mesenchymal cell lines also expressed high levels of the EMT transcription factor *Zeb2*, together with reduced expression of the epithelial *Zo1*, *Sox17*, and *Ghrl2* genes (**Fig. 3D**). Mesenchymal cell lines showed increased motility in a wound closure assay *in vitro* compared to CDH1-positive cell lines, independent of tumor genotype (**Fig. S6F**). Several genes of canonical and non-canonical WNT signaling were upregulated in mesenchymal PDAC cells including *Wnt5a*, *Wnt10b*, while a negative regulator of WNT signaling, *Rnf43* (Wu et al., 2011), was nearly undetectable (**Fig. 3D**). As anticipated, the expression of CDH1, a hallmark feature of epithelial lineage, negatively correlated with expression of *Zeb2, Wnt5a, Wnt10a* and secreted *Tgfb1* to the media, whereas epithelial biomarkers *Zo1*, *Sox17*, cholesterol pathway *Tm7sf2*, *Dhcr24* and *Ebp*, and a negative regulator of WNT signaling *Rnf43* correlated with CDH1 expression (**Fig. 3E, S6G**). In addition to mRNA expression differences, we specifically probed the signaling activity of the TGFβ pathway reflected by levels of phosphorylated SMAD3, a canonical TGFβ pathway effector (Kretzschmar et al., 1999), and found a strong negative correlation with CDH1 and cholesterol pathway genes (**Fig. 3E, S6G**).

Notably, TGFβ1 is a major regulator of stromal activation in PDAC (Franco-Barraza et al., 2017), and increased secretion of this protein could explain the increased proliferation of stromal CAFs observed in *KPPCN* premalignant lesions (**Fig 2G, H**). An additional common regulator of PDAC stromal activation is Sonic hedgehog (Shh) (Rhim et al., 2014). Comparison of Shh secretion in cell culture supernatants produced by PDAC cells derived from *KPPC* and *KPPCN* pancreatic tumors (**Fig. S7A**) indicated no difference in Shh secretion. This suggested NSDHL deficiency alone, in the absence of stromal cells or TGFβ1, was not inducing tumoral Shh secretion *in vitro*.

### Cholesterol deprivation activates TGFβ signaling

To establish a functional linkage between *Nsdhl* deficiency, decrease in the expression of cholesterol biosynthesis pathway enzymes, and EMT, we first established that total cellular cholesterol of *KPPCN* cell lines was approximately 25% lower than in *KPPC* cells when grown in cell culture medium containing lipids (with fetal bovine serum, FBS) indicating that cellular uptake of exogenous lipids and cholesterol could not fully compensate for the reduced endogenous cholesterol biosynthesis. These differences were further exaggerated in cultures supplemented with lipid-depleted serum (LDS, **Fig. 4A**). We proceeded next to investigate a possible causal link between cellular cholesterol levels and activation of the TGFβ pathway in PDAC cells. To induce a decrease in total cellular cholesterol comparable to *KPPCN* cell lines, we cultured *KPPC* PDAC cell lines (KPC3 and KPC634) for 48 hours in LDS medium with non-toxic concentrations of the 3-hydroxy-3-methylglutaryl-coenzyme A reductase inhibitor, compactin(Brown et al., 1978) (**Fig. 4B**). Strikingly, *KPPC* cell lines grown in LDS with compactin in the absence of added TGFβ1 showed levels of phosphorylated SMAD2/3 comparable to a 30-minutes pulse with TGFβ1 in cholesterol-replete cells (**Fig. 4C, D**). The increased levels of pSMAD2/3 induced by LDS+compactin were eliminated by SB431542, a selective inhibitor of the type I TGFβ receptor tyrosine kinase 1 (TGFBR1) and the homologous activin receptors ACVR1B and ACVR1C (**Fig. 4C**). Silencing of TGFBR1, but not of ACVR1B and ACVR1C, with siRNA demonstrated abrogation of pSMAD2/3 in cholesterol-depleted KPC3, indicating TGFβ family receptors predominated in the activation of SMAD2/3 in these cells (**Fig. 4E, S6H**).

**Figure 4.**
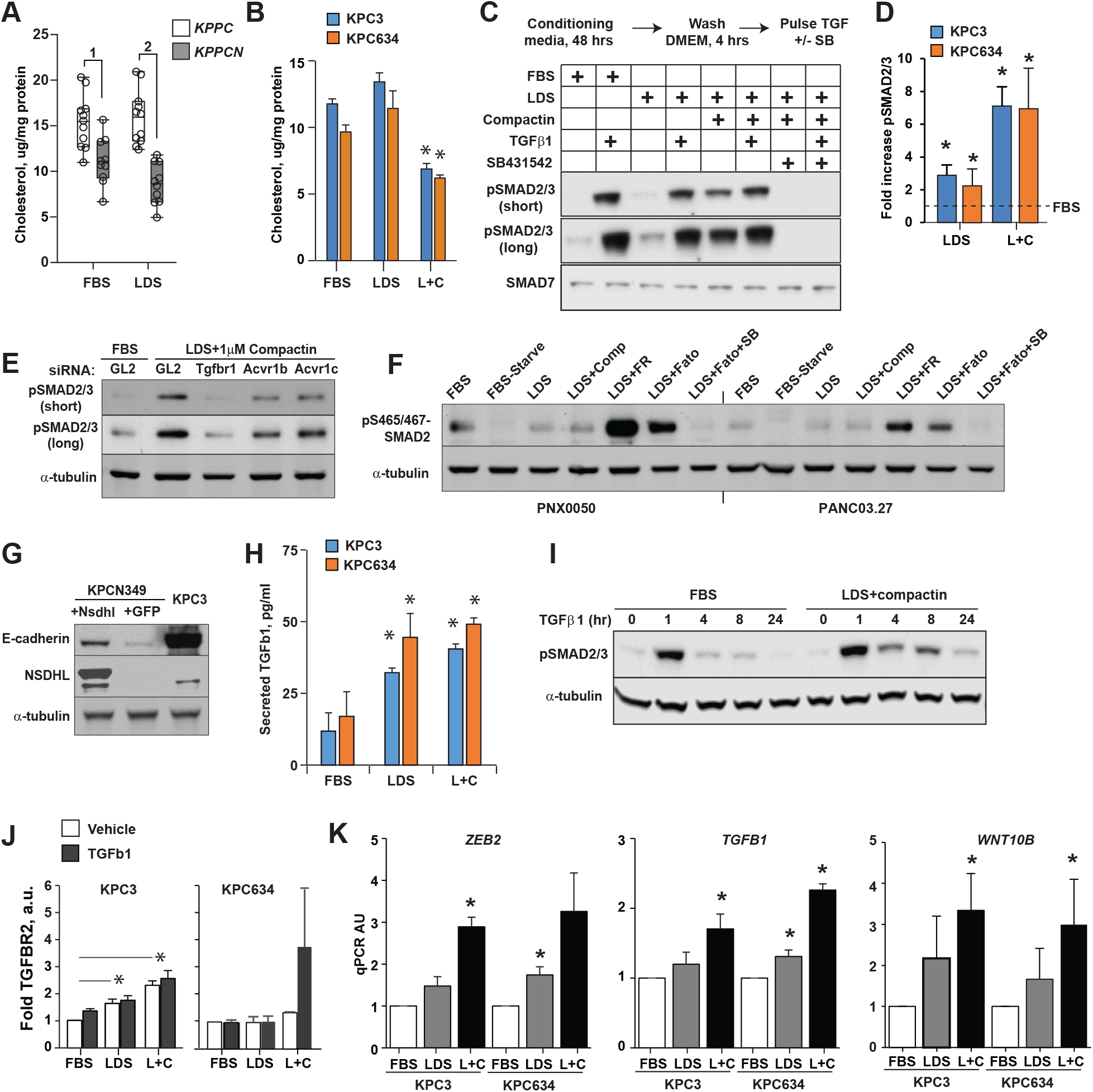
Cholesterol deprivation activates TGFβ pathway signaling in PDAC cells. (**A**) Cholesterol level in *KPPC* (n=10) and *KPPCN* (n=10) clones grown for 48 hours in FBS or LDS media; (1) *p*=0.004; (2) *p*=0.0001, Wilcoxon test. (**B**) Cholesterol levels in PDAC cells conditioned for 48 hours as indicated; L+C, 5%LDS with 1 µM Compactin. *, *p*<0.05, two-way Student t-test. Cholesterol depletion increased SMAD2/3 phosphorylation. (**C**) Representative Western blot of phosphorylated pSMAD2 (Ser465/467) and pSmad3 (Ser423/425) in KPC3 cells cultured for 48 hours in fetal bovine serum or lipid depleted serum (LDS) in DMEM, followed by incubation in serum-free DMEM for 4 hours. Indicated samples were treated with TGFβ1 at 10 ng/ml for 30 minutes, or SB431542 at 25 µM for 1 hour. (**D**) Summary results of cholesterol depletion on levels of phosphorylated pSMAD2(Ser465/467) and pSmad3 (Ser423/425). Shown are results from 3 independent experiments normalized to α-tubulin. (**E**) Silencing of TGFBR1, ACVR1B and ACVR1C with siRNA abrogated SMAD2/3 phosphorylation in KPC3 cells preconditioned for 48 hours in LDS with compactin. (**F**) Effect of cholesterol depletion on expression of pSMAD2/3 in human PDAC cell lines. Cells were conditioned for 48 hours in the presence of 1 µM Compactin (Comp), 10 µM FR171456 (NSDHL inhibitor, FR), 20 µM Fatostatin (SREBP1/2 inhibitor, Fato), or 25 µM SB431542 (TGFBR1 inhibitor, SB). (**G**) Expression of E-cadherin in KPCN349 cells modified to express full length *Nsdhl*. Cells modified with lentiviral vector to express GFP served as control. (**H**) Increased TGFβ1 in supernatants of KPC3 cells conditioned as in (A). (**I**) Persistence in SMAD2/3 phosphorylation upon TGFβ1 treatment. KPC3 cells were conditioned in FBS or LDS+compactin with for 48 hours, pulsed with 2.5 ng/ml of TGFβ1 for 60 min, washed and lysed at the times indicated. Levels of pSMAD2 and tubulin as a loading control were assayed by Western blotting. (**J**) Quantified expression of TGFBR2 in KPC3 and KPC634 cells conditioned in FBS, LDS, or LDS+compactin 1 µM (L+C). (**K**) Expression of *Zeb2*, *Tgfb1* and *Wnt10b* mRNA as assessed by qRT-PCR in KPC3 and KPC634 PDAC cells cultured for 48 hours in media supplemented with FBS, LDS or LDS+compactin (1 µM); *, *p*<0.05, two-way Student t-test.

The induction of pSMAD2/3 by an agent targeting the cholesterol biosynthesis pathway was not specific to compactin or to murine PDAC models, as inhibiting endogenous cholesterol biosynthesis with FR171456, a selective NSDHL inhibitor (Helliwell et al., 2015), or fatostatin, an SREBP inhibitor (Chen et al., 2018; Talebi et al., 2018), similarly upregulated pSMAD2/3 in two human PDAC cell lines (**Fig. 4F**). In addition to increasing SMAD2/3 phosphorylation, cholesterol starvation induced formation of actin stress fibers (**Fig. S6G**), which represent an EMT-associated, SMAD-independent response to TGFβ pathway activation, mediated by the CDC42 and RhoA GTPases (Edlund et al., 2002). Importantly, restoration of cholesterol biosynthetic competency in mesenchymal/basal-like KPCN349 PDAC cells, via lentiviral re-expression of *Nsdhl* cDNA, partially restored CDH1 expression, indicating a direct effect on epithelial differentiation (**Fig. 4G**).

The fact that inhibitors of TGFβ receptors blocked induction of pSMAD2/3 by cholesterol depletion suggested a mechanism by which cholesterol reduction activates this signaling cascade. We had noted that mesenchymal PDAC cell lines expressed higher levels of *Tgfb1* mRNA (**Fig. 3D**). Similarly, KPC3 and KPC634 cells preconditioned for 48 hours in LDS, with or without compactin, secreted higher levels TGFβ1 in culture media (**Fig. 4H**). Several reports have implicated membrane cholesterol as influencing endocytic trafficking of TGFβ receptors, thus affecting TGFβ pathway signaling activity (Di Guglielmo et al., 2003; Miller et al., 2018). We conducted TGFβ1 pulse-chase experiments in which PDAC cells were pre-conditioned in cholesterol-replete or cholesterol-poor cultures and pulsed using low dose, 2.5 ng/ml, TGFβ1. Activation of SMAD2/3 was then followed in the absence of added TGFβ1. These experiments demonstrated markedly prolonged phosphorylation of SMAD2/3 in cholesterol depleted PDAC cells (**Fig. 4I**). Signaling activity of TGFβ receptors is restricted by their endocytic trafficking to late endosomes and lysosomes (Miller et al., 2018), while protracted SMAD2/3 phosphorylation is associated with increased constitutive levels of TGFβ receptor as we determined in cholesterol depleted cells (**Fig. 4J**). We further determined that cholesterol starvation of epithelial PDAC cells increased the expression level of the mesenchymal biomarkers, such as *Tgf*β*1, Zeb2* and *Wnt10b* (**Fig. 4K**) defined by transriptome analyses of PDAC cell lines (**Fig 3D**), implying the switch between subclasses is dynamic and reversible.

To determine whether *in vitro* observations of cholesterol-dependent modulation of TGFβ signaling activity correlated with the effects on PDAC development observed in the KPC mouse model, we evaluated activation of this pathway in murine tumor tissues. In agreement, NSDHL-deficient premalignant pancreatic lesions demonstrated more activated pSMAD2, than did lesions in *KPC* and *KPPC* mice (**Fig. S7D**).

### Distinct cholesterol metabolic programs define transcriptional subsets of human pancreatic cancer

To determine whether the relationship between the cholesterol biosynthesis and EMT signaling observed in mouse models is also reflected in human PDAC, we compared transcriptional profiles between classic and basal subsets of PDAC using data available for 76 high purity pancreatic adenocarcinoma samples by The Cancer Genome Atlas Research Network (TCGA., 2017). Notably, basal PDAC had significantly increased expression of the gene signatures of TGFβ and EMT signaling, among other hallmark pathways of tumor aggressiveness such as the PI3K-mTOR pathway, mitosis, and hypoxia (**Fig. 5A**). In contrast, the fatty acid and cholesterol metabolism gene signatures (Molecular Signature Database, (Liberzon et al., 2015)) were significantly higher in classic PDAC compared to basal (**Fig. 5B**). In confirmatory analysis, we found that the cholesterol biosynthesis gene signature (Reactome, (Fabregat et al., 2017)) was significantly lower in basal PDAC, compared to classic, from independent cohorts curated by the International Cancer Genome Consortium (ICGC, **Fig. 5C**).

**Figure 5.**
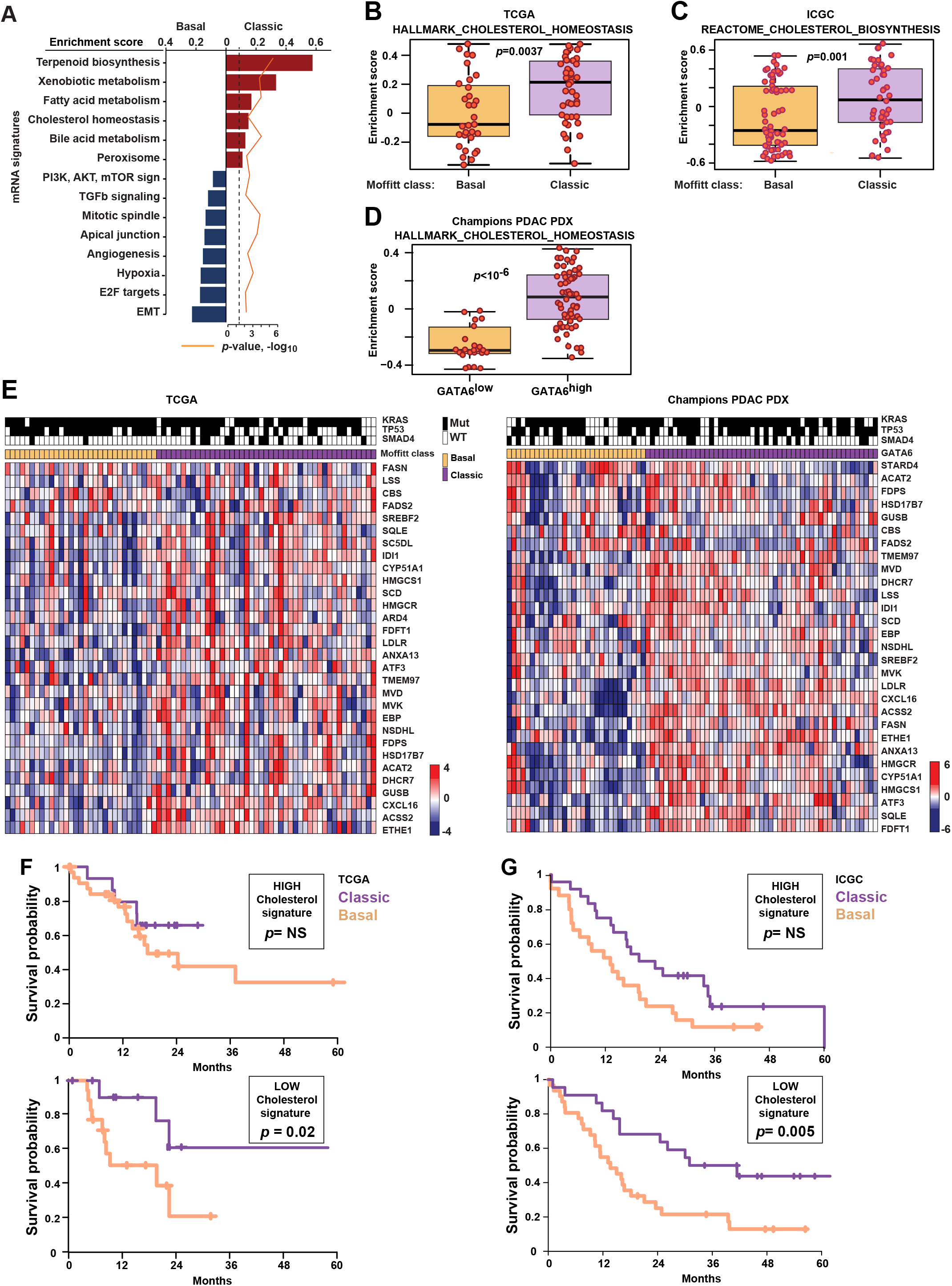
Cholesterol homeostasis regulates classic versus basal differentiation of human PDAC. (**A**) Comparison of mRNA transcriptional signatures between classic and basal PDAC among 76 TCGA cases with estimated tumor cell fraction greater than 30% and *p*-values <0.01 (shown as –log_10_, orange line). (**B-D**) Comparison of cholesterol homeostasis genes expression in human basal and classic PDAC. In **B**, expression of “hallmark_cholesterol_homeostasis” signature (Molecular Signature Database(<u>Liberzon et al., 2015</u>) in 76 TCGA PDAC cases; in **C,** shown are “reactome_cholesterol_biosynthesis” mRNA signatures in 102 PDAC cases from ICGC (GSE50827); and **D** represents “hallmark_cholesterol_homeostasis” mRNA signature expression in 85 patient-derived xenograft samples stratified by human GATA6 mRNA Z-score above or below zero. y-axes illustrate enrichment score comparing basal and classic subtypes of PDAC. Positive and negative scores indicate positive and negative enrichment, respectively. Boxplots represent median (black bar) and range (25-75^th^ percentile) of enrichment scores for individual cases shown as red dots for each sample in that subtype. (**E**) Heat map of normalized expression of representative genes in the “hallmark_cholesterol_homeostasis” signature for TCGA and human PDAC PDX cases. Z-scores calculated for each gene are plotted on a red (higher expression) and blue (low expression) scale. *Top color bar*, subtype of PDAC. *Mutations*, shown as black lines if present. (**F, G**) Kaplan-Meier survival of Moffitt classic and basal PDAC in the TCGA (**F**) and ICGC (**G**) cohorts stratified by positive or negative enrichment for “hallmark_cholesterol_homeostasis” signature. See also Tables S3-4.

Separately, we classified a panel of 85 human PDAC patient-derived xenografts (PDXs) by a validated clinical biomarker(Aung et al., 2018), i.e. high or low *GATA6* mRNA expression, as classic or basal PDAC, respectively. In accord with the TCGA and the ICGC datasets, mRNA analysis of *GATA6*^low^ PDX showed significantly lower expression of hallmark cholesterol homeostasis genes (**Fig. 5D**). The cholesterol homeostasis genes underrepresented in basal PDAC (**Fig. 5E**) included 16 canonical genes of cholesterol biosynthesis, *LDLR* (the principal receptor for exogenous cholesterol uptake (Guillaumond et al., 2015)), and *SREBF2*, a master transcription factor of cholesterol biosynthesis pathway (Hua et al., 1995). These data strongly suggested functional cholesterol metabolic auxotrophy in basal subset of human PDAC. Importantly, the cholesterol homeostasis gene signature independently impacted the overall survival of patients. For these survival analyses, we compared PDAC cases from TCGA and ICGC datasets stratified by their positive or negative cholesterol gene signature enrichment scores (**Fig. 5B, C**). The cases with positive enrichment for cholesterol homeostasis genes had a similar, longer survival, irrespective of the Moffitt basal or classic subclass. Contrastingly, low expression of cholesterol homeostasis genes was associated with dismally shorter survival of the basal PDAC patients compared to the classic variant (**Fig. 5F, G**) thus indicating poorer prognosis in patients with low cholesterol pathway genes expression.

## Discussion

Pancreatic cancer remains one of the most lethal human cancers. EMT and uncontrolled metastases are the primary cause of mortality in PDAC, and particularly associated with the basal subtype (Aiello et al., 2018; Rhim et al., 2012). This study provides evidence for a direct role of cholesterol metabolism in induction of EMT and basal PDAC development. In the mouse model, disruption of cholesterol biosynthesis consistently decreased tumor differentiation in Kras-dependent pancreatic tumorigenesis, in a phenotype we defined as cell-intrinsic. Mechanistically, this arose from upregulated expression of TGFβ1, which acted in an autocrine manner through SMAD2/3 to promote EMT in PDAC cancer cells, and in a paracrine manner to activate CAFs surrounding the cancer cells, which are known for their increased TGFβ production(Kojima et al., 2010). We also showed that starvation for cholesterol in cells with intact NSDHL separately resulted in similar induction of EMT, based on activation of TGFβ1 signaling. Finally, in human PDAC, we established a relationship between a signature indicative of reduced cholesterol homeostasis and poor survival, specifically in the basal subset of PDAC tumors, which constitute ∼25% of PDAC overall. Given the known linkages between diet, obesity, and the risk and prognosis of aggressive cancers (Golemis et al., 2018), there has long been interest in the relationship between cholesterol and PDAC pathogenesis (Huang et al., 2017). The complex signaling mechanism revealed in this study offers one explanation as to why such relationships have been difficult to establish, and for the poor clinical outcomes in trials of statins for PDAC (Hong et al., 2014). Experimental models and clinical trials of statins aimed to define the role of cholesterol metabolism in disease risk and treatment have yielded complicated and sometimes conflicting results (Chen et al., 2019; Huang et al., 2017). Importantly, our data provides a mechanistic framework for interpretation of contradictions arising from some of these experiments. For instance, our results raise the possibility that potential beneficial effects cholesterol restriction could be confined to cancers with functional P53: a relatively small cohort. In the broader population of PDAC patients, cholesterol lowering interventions may have inadvertently favored the selection of poorly differentiated cancer cells that have undergone EMT and contributed to resistance and worse outcomes. Further, based on our transcriptome analyses (**Fig. 5**), the effects of statins might only be relevant to the classic rather than the basal subset of PDACs. In the latter, lowering cholesterol may be detrimental by further promoting TGFβ signaling. Our retrospective analyses may be of interest for further prospective validation in the clinic and for the applicability of cholesterol suppression as a therapeutic modality in PDAC. As an example, our results may explain why statin use has been associated with modestly improved overall survival in patients with existing pancreatic tumors, but where the reductions in level of serum cholesterol did not correlate with survival in individual patients (Huang et al., 2017).

Genetic ablation of *Nsdhl* provides a useful model for dissection of cholesterol signaling, as loss of its activity in catalyzing an irreversible step of oxidative C4-demethylation is more selective to cholesterol instead of mevalonate pathway inhibitors having pleiotropic effects on farnesylation and geranylation-based signaling (Deng et al., 2018). While inactivating mutations in *NSDHL* or other cholesterol pathway genes are extremely rare in pancreatic cancer (*<u>cbioportal.org</u>*), low *NSDHL* mRNA expression was evident in the majority of “basal” PDAC (**Fig. 5**). Conversely, a signature of higher expression of multiple additional cholesterol pathway genes is a feature of classic PDAC. An intriguing feature of our study is the fact that non-equivalent results were obtained when *Nsdhl* loss was accompanied by heterozygous versus homozygous loss of *Trp53*. *Nsdhl* inactivation in the context of the heterozygous loss of *Trp53* dramatically delayed malignant progression of the precursor lesions and nearly completely abrogated PDAC development (**Fig. 1A**). These results indicate that in the context of partial retention of *Trp53* tumor suppressor activity, cholesterol pathway proficiency is critical for the continuous progression from ADM to PanINs, and ultimately to the classic (glandular/classic) variant of PDAC associated with an activating *Kras* mutation (**Fig. 1B**). Conversely, the linear ADM-PanIN-classic PDAC progression was impaired in *KPPCN* mice with complete deletion of *Trp53*, and hence with reduced restriction on proliferation (**Fig. 1C,** (Morton et al., 2010)). Despite this reduced ability to induce classic PDAC, all *KPPCN* mice died of fulminant basal PDAC originating from rare microscopic foci, expressing elevated TGFβ1 and surrounded by highly activated yet non-canonical CAFs.

The mechanisms by which cholesterol deprivation stimulates TGFβ signaling and EMT require further analysis. Among candidate mechanisms, depletion of the cellular membrane cholesterol pools has been shown to increase membrane fluidity promoting EMT (Zhao et al., 2016). In line with our findings, genetic studies in breast cancer mouse model have demonstrated induction of the EMT by increased levels of 27-hydroxycholestrol, an established LXR agonist (Nelson et al., 2013). Based on these data, reduced biosynthesis of cholesterol could be nominated as an EMT-promoting metabolic switch by reinforcing TGFβ1 pathway signaling in the absence of P53. In human PDAC, we hypothesize that a nutrient- and oxygen-restricted tumor microenvironment may lower the levels of cholesterol in early stage PanINs, activating TGFβ signaling and supporting emergence of basal PDAC. It is possible that basal PDAC are ultimately selected to bypass a strong anti-proliferative and pro-apoptotic effects of TGFβ signaling elicited through SMADs and P53 interactions (Cordenonsi et al., 2003) induced in cholesterol-poor tumors. Therefore, P53 is nearly universally mutates in basal PDAC (**Fig. 5E**) where it exerts a dominant negative effect to overcome the KRAS-induced senescence and promote PDAC metastases(Hingorani et al., 2005; Morton et al., 2010). Our use of conditional *Trp53^f/f^* mice instead of P53 mutant allowed us to functionally probe the complex interactions between P53, TGFβ pathway and cholesterol metabolism during KRAS-induced pancreatic carcinogenesis. Some data suggests that a role of cholesterol in regulating TGFβ pathway signaling may also be relevant to other human pathologies in which cholesterol depletion or accumulation takes place. One interesting example is in formation of atherosclerotic plaques, where progressive cholesterol accumulation abolishes TGFβ signaling and SMAD3 phosphorylation in smooth muscle cells (Chen et al., 2016).

## Author contributions

L.G.C., S.P., and I.A. designed research; L.G.C., S.P., D.R., A.K., T.R.H, R.F., J.F.-B., N.S., E.N., E.H., K.Q.C., I.S. and R.A.M. performed research; R.A.M., R.C. and A.M.O. contributed new reagents/analytical tools; L.G.C., S.P., E.H., E.C. and I.A. analyzed data; L.G.C., E.A.G, E.C. and I.A. wrote the paper.

## Acknowledgments

We are grateful to Catherine Reiner and the FCCC Immune Monitoring Facility for technical assistance with histological analyses and immunohistochemistry experiments. **Grant Support:** This work was supported by NIH core grant CA-06927, by the Pew Charitable Fund, and by a generous gift from Mrs. Concetta Greenberg to the M&C Greenberg Pancreatic Cancer Institute at Fox Chase Cancer Center. Some of the authors were supported by NIH R01 CA188430, K22 CA160725, R21 CA164205, R21 CA231252 (I.A., E.C.), a career development award from Genentech; by Tobacco Settlement funding from the State of Pennsylvania (I.A., E.C.), and by a grant from the Bucks County Board of Associates (L.G.C., I.A.) as well as by NIH R01 CA63366 (E.A.G.); R01CA113451 (E.C.); WX81XWH-15-1-0170 (E.C., R.F., J.F.-B.), R01 CA232256 (E.C., J.F.-B.), T32CA009035 (R.F., L.G.C.), R01 HD065800 (A.M.O.) and by the Program of Competitive Growth of Kazan Federal University (L.G.C.); cloning experiments were supported by the grant from Russian Scientific Foundation (project 15-15-20032) to I.A.

## METHODS

Detailed methods are provided in the online version of this paper and include the following:

- KEY RESOURCES TABLE
- CONTACT FOR REAGENT AND RESOURCE SHARING
- EXPERIMENTAL MODEL AND SUBJECT DETAILS
  1. Mouse strains
  2. Cell lines
- METHOD DETAILS
  1. PCR genotyping
  2. Induction of acute pancreatitis in mice with caerulein
  3. Micro-magnetic resonance imaging (mMRI)
  4. Orthotopic reimplantations
  5. Immunohistochemistry
  6. Histology images analysis
  7. Lesion and tumor grading criteria
  8. Simultaneous multi-channel immunofluorescent (SMI) labeling of formalin-fixed paraffin-embedded mouse pancreatic tumors
  9. mNSDHL expression in KPCN cells
  10. WB analysis of protein expression levels in cell lines and tissues
  11. Quantitative RT-PCR
  12. Clustering analysis
  13. RNAseq and its bioinformatical analysis
  14. Wound healing assay
  15. TGFβ ELISA
  16. Confocal image acquisition
  17. Cholesterol measurement
  18. siRNA transfection
  19. Shh ELISA
  20. Phalloidin staining of stress fibers
- STATISTICAL ANALYSIS
- DATA AND SOFTWARE AVAILABILITY

### KEY RESOURCES TABLE

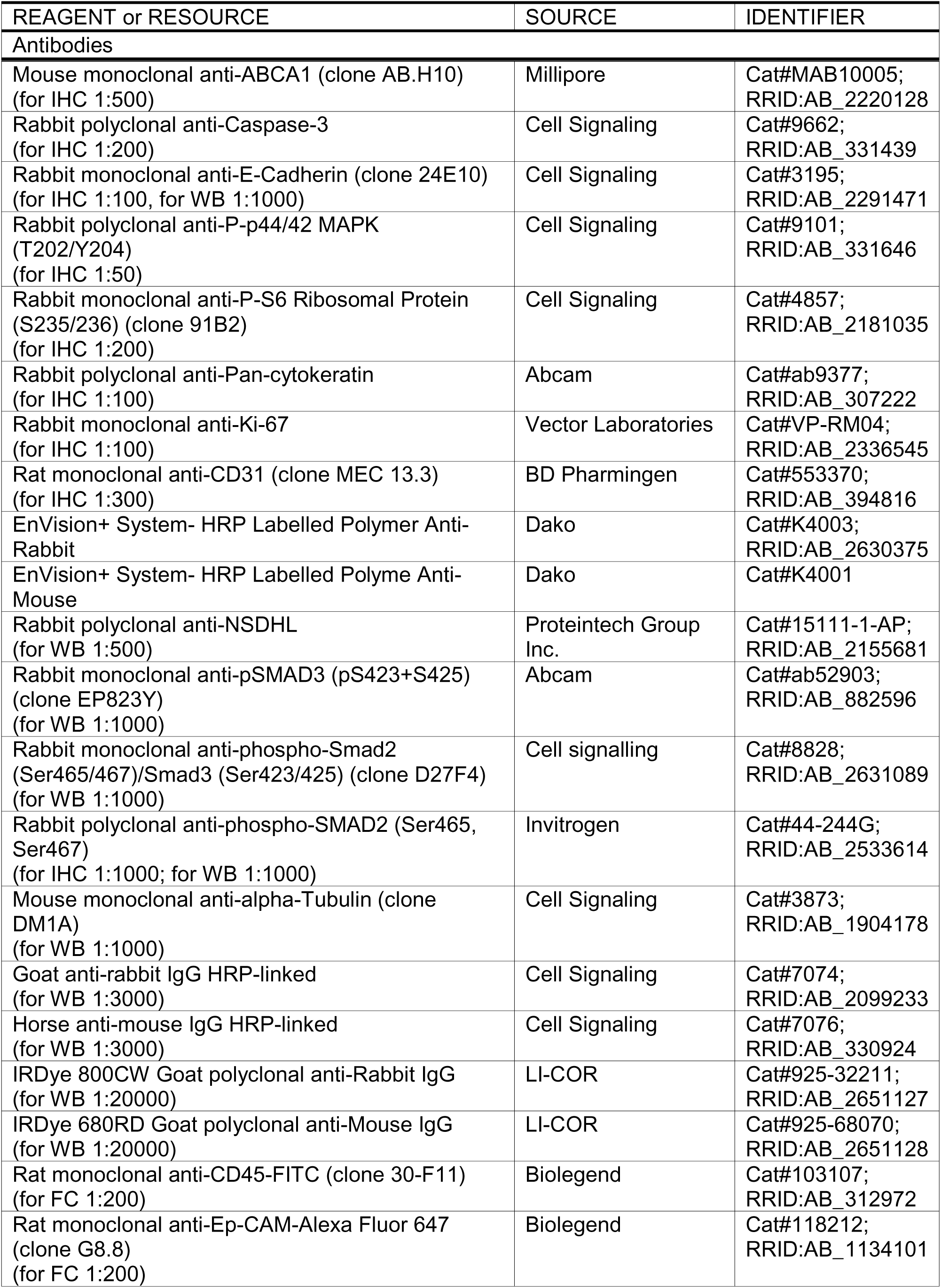
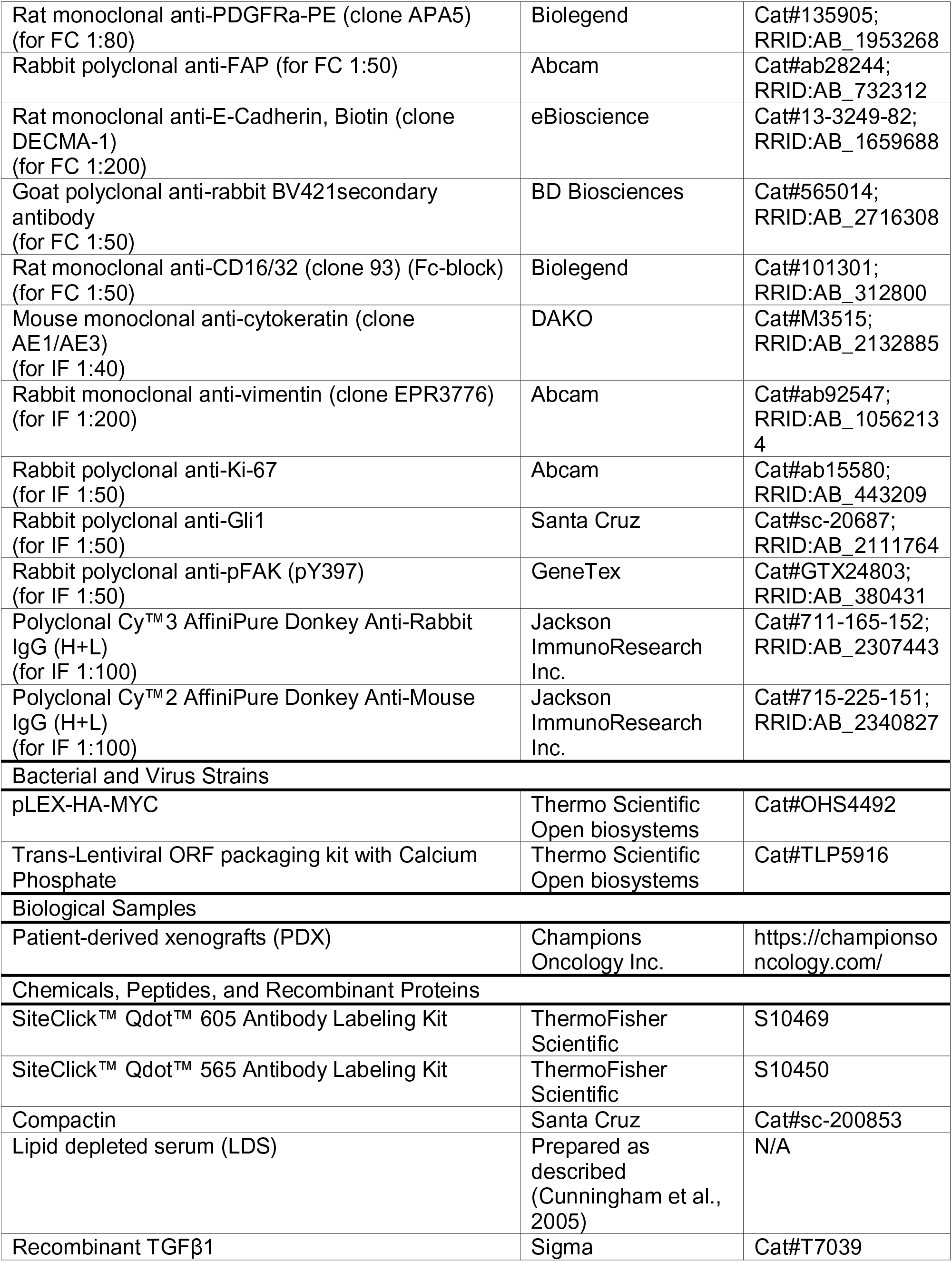
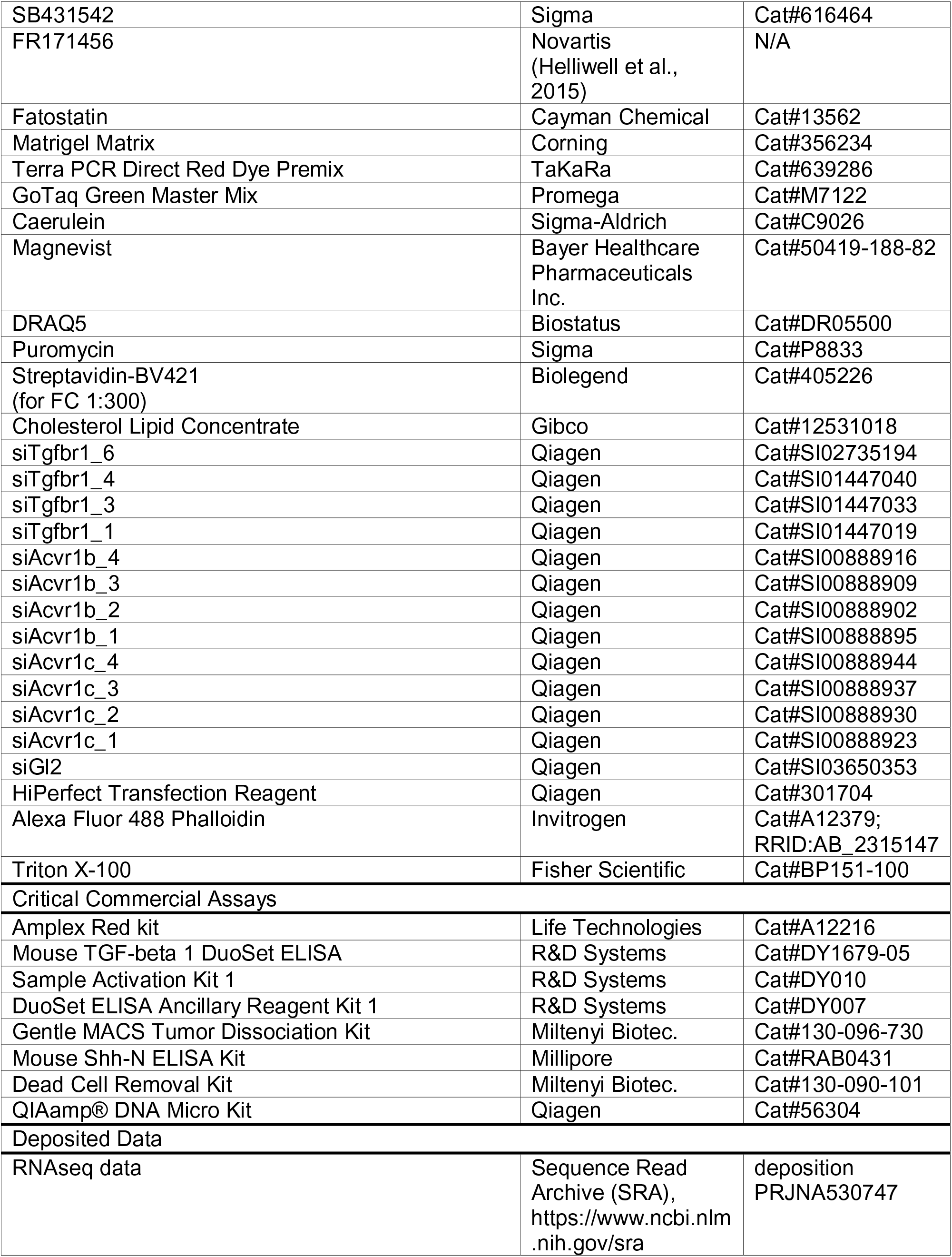
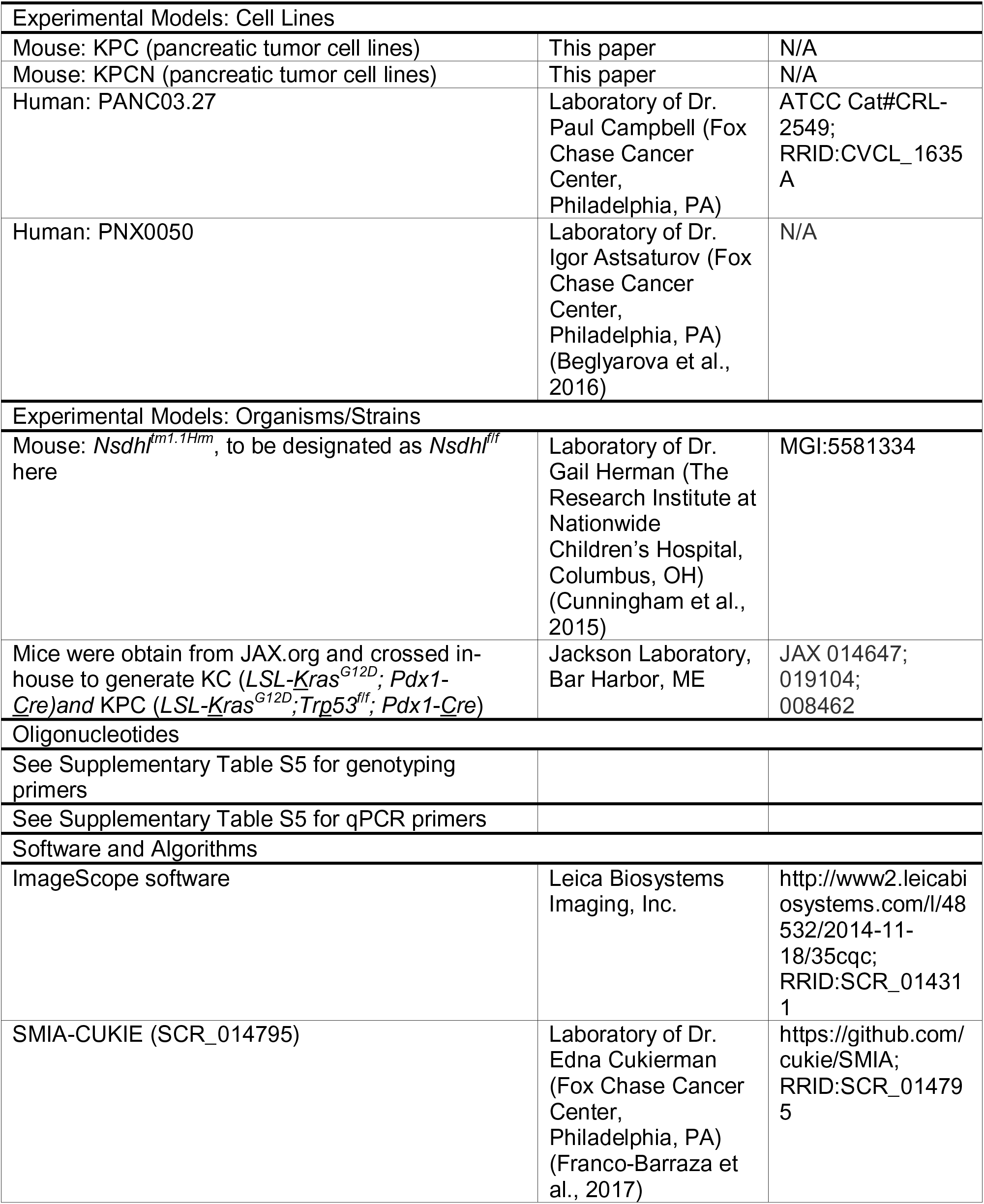
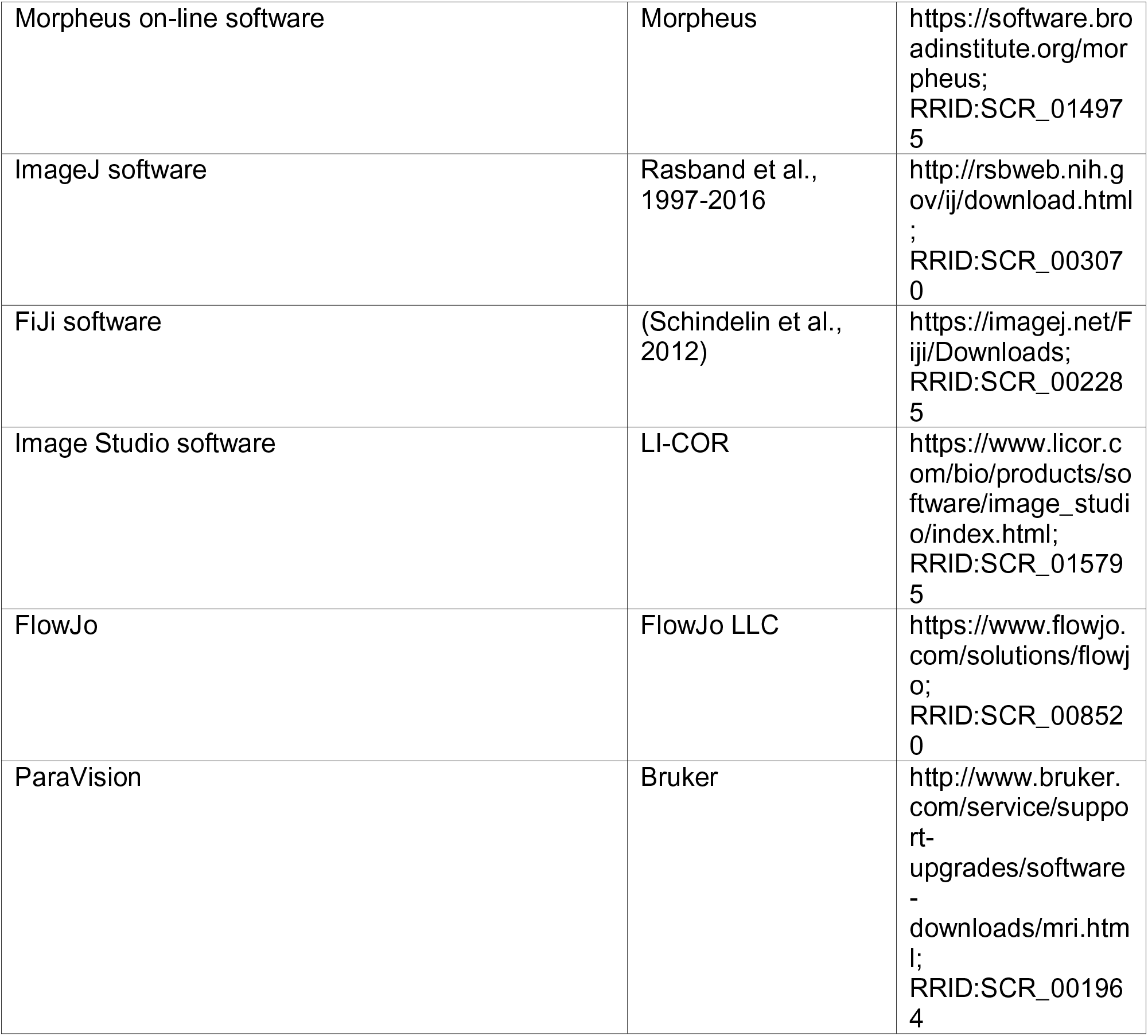

### CONTACT FOR REAGENT AND RESOURCE SHARING

Further information and requests for resources and reagents should be directed to and will be fulfilled by the Lead Contact, Igor Astsaturov.

### EXPERIMENTAL MODEL AND SUBJECT DETAILS

#### Mouse models

Mice carrying the conditional knockout allele of *Nsdhl* (Cunningham et al., 2015) were kindly provided by Dr. Gail Herman (The Research Institute at Nationwide Children’s Hospital, Columbus, OH). These mice (official name: *Nsdhl^tm1.1Hrm^*, MGI:5581334, to be designated as *Nsdhl^f/f^*here) are congenic on a C57BL/6J background. KPC (*LSL-<u>K</u>ras^G12D^;Tr<u>p</u>53^f/f^; Pdx1*-*<u>C</u>re*) and KC (*LSL-<u>K</u>ras^G12D^; Pdx1*-*<u>C</u>re)* mice were kindly provided by Dr. Kerry Campbell (Fox Chase Cancer Center, Philadelphia, PA) and those mice are congenic on a C57BL/6J background as well. Mice were bred to obtain the desired genotype with *Nsdhl* depletion in normal pancreas (*Nsdhl^f/f^; Pdx1-Cre*, to be designated as *Nsdhl*^Δ^*P^anc^* here) and *Nsdhl* depletion in pancreas of KPC mice with different status of p53 expression (KPC mice carrying *Trp53^f/f^* or *Trp53^f/+^*, or KC mice). All mice were bred and kept under defined-flora pathogen-free conditions at the AAALAC-approved Animal Facility of the Fox Chase Cancer Center, Philadelphia, PA. Mice of both genders, equally distributed, were used for experiments. Tumor bearing mice were observed once weekly until signs of sickness appeared or animals showed distress or weight loss of more than 10%, per the local Institutional Animal Care and Use Committee (IACUC) guidelines.

#### Cell Lines

Mouse pancreatic cancer cell lines (KPC and KPCN) were derived from mouse pancreatic tumors by tumor dissociation and subsequent fluorescence-activating cell sorting (FACS). Tumor dissociation was performed with the use of the Gentle MACS Tumor Dissociation Kit (Miltenyi Biotec, Order No. 130-096-730) according to the manufacturer’s instructions. Briefly, each tumor was isolated from the animal in a sterile environment and washed in PBS. A 1mm^3^ piece was taken for genotyping and a larger piece was taken for histopathology analysis. The rest of the tumor tissue was placed in the dissociation enzyme mix and minced quickly to get pieces ∼2mm^3^ in size. Then the enzyme-tissue mixture was transferred into the gentleMACS C tube and incubated at 37°C with constant rotation for 40 minutes. After that the tissue was further mechanically processed by the gentleMACS Dissociator. The single-cell suspension was obtained by passing the tissue mixture through the 70µm cell strainer. Dead cells were subsequently removed by the Dead Cell Removal Kit (Miltenyi Biotec, Order No. 130-090-101).

In order to separate cells of different origin we further stained the cell suspension with antibodies against CD45 (Biolegend #103107; 1:200), FAP (Abcam #ab28244; 1:50), EPDACAM (Biolegend #118212; 1:200) and PDGFRa (CD140a) (Biolegend #135905; 1:80). To prevent antibodies from binding to Fc-receptors we treated the cell suspension with Fc-block (Biolegend #101301; 1:50) prior using other antibodies. To detect cells labeled with FAP we used BV421 anti-rabbit secondary antibody (BD Biosciences #565014; 1:50). Live cells were selected based on Propidium Iodide staining (Biolegend #421301). Fluorescence detection and sorting were performed with BD FACS Aria II flow cytometer. Sorted cells were propagated for first two passages in enriched media (RPMI-1640 supplemented with 15% v/v FBS, 2mM L-glutamine, 100µg/ml Penicillin/Streptomycin, 20ng/ml EGF, 25µg/ml Insulin, NEAA 1x, 1mM Na-Pyruvate and 2µg/ml Hydrocortizone) and for subsequent passages in DMEM supplemented with 10% v/v FBS and 2mM L-glutamine with 100µg/ml Penicillin/Streptomycin. Cells were used at passages 3-5.

Human pancreatic cancer cell line PANC03.27 (#CRL-2549, ATCC) was kindly provided by Dr. Paul Campbell (Fox Chase Cancer Center, Philadelphia, PA).

Human pancreatic cancer cell line PNX0050 was derived as described earlier (Beglyarova et al., 2016).

### METHOD DETAILS

#### PCR genotyping

Small (1-2mm) pieces from mouse tails or mouse pancreata were used for genotyping. For cell line genotyping we isolated genomic DNA with QIAamp® DNA Micro Kit (#56304, Qiagen). Primers for genotyping are listed in Supplementary Table S5. The PCR reaction was performed using Terra PCR Direct Red Dye Premix (#639286, TaKaRa) for LSL-KRas construct detection and GoTaq Green Master Mix (#M7122, Promega) for all other constructs.

#### Induction of acute pancreatitis in mice with caerulein

Acute pancreatitis in mice was induced by caerulein (#C9026, Sigma-Aldrich) treatment as previously described (Morris et al., 2010). Briefly, at day “-1”, mice were injected with 50µg/kg of caerulein i/p every hour for 6 hours (6 injections total). In 24 hours (day “0”) the course of injections was repeated. Control mice were injected with saline. At days “2”, “5”, “7” pancreatic tissues were collected and immediately fixed in formalin. Fixed tissues were embedded in paraffin and stained with H&E and Ki-67 for further histological analysis.

#### Micro-magnetic resonance imaging (mMRI)

mMRI was performed on 7-8 week old KPPC and KPPCN mice following the standard procedure of Small Animal Imaging Facility (Dr. Harvey Hensley, Fox Chase Cancer Center, Philadelphia, PA). Briefly, animals were imaged in a 7 Tesla vertical wide-bore magnet, using a Bruker DRX 300 spectrometer (Billerica, MA) with a micro-imaging accessory, which included a micro 2.5 gradient set, a 30 cm radiofrequency coil, and Paravision software (Bruker, Billerica, MA). In order to increase the contrast of the imaging, Magnevist (#50419-188-82, Bayer Healthcare Pharmaceuticals Inc.) was diluted in sterile PBS to achieve 46.9mg/ml concentration and injected subcutaneously into the shoulder region, 200µl/mouse, immediately preceding the scan. Prior to Magnevist injection, and throughout the whole imaging process, mice were anesthetized with a mixture of oxygen and isoflurane (2-3%) according to the local IACUC guidelines. Scout scans were performed in the axial and sagittal orientations, permitting us to accurately prescribe an oblique data set in coronal and sagittal orientations that included the organs of interest. A two dimensional spin echo pulse sequence was employed with echo time 15msec, repetition time 630msec, field of view=2.56cm, acquisition matrix=256×256, slice thickness=0.75mm, 2 averages, scan time=5 minutes. Fat suppression (standard on Bruker DRX systems) was used for all scans.

#### Orthotopic reimplantations

Orthotopic reimplantations of KPC and KPCN tumor cells in mouse pancreas were performed as described by Kim *et al*. (Kim et al., 2009). Total 10^6^ cells per 50µl of 30%/70% Matrigel/PBS mixture were injected into the pancreatic head region of syngeneic C57BL/6J mice. Anesthetics and analgesics were used according to the local IACUC guidelines. Tumor bearing mice were observed twice weekly until signs of sickness appeared or animals showed distress or weight loss of more than 10%, per the local IACUC guidelines.

#### Immunohistochemistry

For immunohistochemistry, formalin-fixed pancreatic tissue was embedded in paraffin and stained with indicated antibodies diluted per the manufacturer’s instructions (listed in Key Resources Table). Antibody binding was visualized using the Liquid DAB+ Substrate Chromogen System (Dako). Samples were counterstained for 1 minute with hematoxylin. Collagen deposition and organization were directly visualized by the standard Masson’s Trichrome Stain.

#### Histology images analysis

Slides were scanned by an Aperio ScanScope CS scanner (Aperio) and selected regions of interest were outlined manually. The surface area taken by different lesions, the expression levels of ABCA1, pS-S6, cleaved Caspase-3, Ki-67, pErk, CDH1, CD31, pSMAD2 and trichrome positive area were measured using the ImageScope software (Leica Biosystems Imaging, Inc.).

#### Lesion and tumor grading criteria

Normal tissue, acinar-to-ductal metaplasia (ADM), pancreatic intraepithelial neoplasia (PanINs) and pancreatic ductal adenocarcinoma (PDAC) were determined according to the standard classification (Hruban et al., 2006). For histopathological scoring, tumors were classified using the standard pathological grading scheme into either well differentiated (grade 1), moderately differentiated (grade 2), poorly differentiated (grade 3) or undifferentiated (which includes sarcomatoid) (grade 4).

#### Multi-channel immunofluorescent labeling of mouse pancreatic tumors

For vimentin, cyto-keratin, Gli1 and pFAK immunofluorescent co-staining of formalin-fixed paraffin-embedded mouse pancreatic tumors, we utilized the simultaneous multi-channel immunofluorescent (SMI) labeling approach described earlier (Franco-Barraza et al., 2017). Briefly, Gli1 and pFAK primary antibodies were conjugated with Q-dot probes by using antibody Q-dot labeling kits (#S10469 and #S10450, ThermoFisher Scientific). Slides were deparaffinized and stained with Q-dot pre-labeled antibodies (Franco-Barraza et al., 2017). To detect epithelial/tumoral locations and mesenchymal (stromal) components slides were further stained with the mouse monoclonal “cocktail” of anti-pan-cytokeratin (clones AE1/AE3, DAKO) and anti-vimentin (EPR3776, Abcam) antibodies with subsequent staining with secondary donkey anti-mouse Cy2 and donkey anti-rabbit Cy3 antibodies. Nuclei were stained using DR (1:10^5^, #DR50050, Biostatus) (Franco-Barraza et al., 2017). Images were collected using Caliper’s multispectral imaging system (Perkin Elmer) and analyzed with SMIA-CUKIE (SCR_014795) software, as described earlier (Franco-Barraza et al., 2017).

#### WB analysis of protein expression levels in cell lines and tissues

For Western blot analysis, the piece of tissue sample or cultured cells were homogenized in RIPA buffer (#24928, Santa Cruz) with phosphatase and protease inhibitors (#1862495, #1861278, Thermo Scientific) on ice and cleared then by centrifugation. The protein concentration was measured with Pierce BCA Protein Assay Kit (#23225, Thermo Scientific). Proteins were separated on the 4-12% Bis-Tris Protein gels (Invitrogen) and then horizontally transferred to the Immobilon-FL PVDF membrane (#IPFL00010, Millipore). Primary and secondary antibodies were used in concentrations indicated in the Key Resources Table according to manufacturer’s instructions. The density of obtained bands was quantified with Image Studio software (LI-COR).

#### Quantitative RT-PCR

For evaluation of target gene expression, total RNA was extracted using the RNeasy Mini Kit (#74104, Qiagen). RNA was reverse transcribed (RT) using Moloney murine leukemia virus (MMLV) reverse transcriptase (#28025013, Ambion) and a mixture of anchored oligo-dT and random decamers (IDT). Two reverse-transcription reactions were performed for each sample using either 100 or 25ng of input RNA in a final volume of 50µl. Taqman or SYBR Green assays were used (see Supplementary Table S5) in combination with Life Technologies Universal Master mixes and run on a 7900 HT sequence detection system (Life Technologies). Cycling conditions were 95°C, 15 minutes, followed by 40 (two-step) cycles (95°C, 15s; 60°C, 60s). Ct (cycle threshold) values were converted to quantities (in arbitrary units) using a standard curve (five points, four fold dilutions) established with a calibrator sample.

#### Clustering analysis

Hierarchical clustering analysis of KPC and KPCN cell lines was performed based on data obtained from qPCR and WB results. The expression level of each gene (for qPCR) or the level of each protein in total cell lysates (for WB) for all analyzed cell lines was first normalized by converting them to a percentage, wherein the highest level of expression for each gene was set to 100%. The obtained results were further clustered using Morpheus on-line software (Morpheus, https://software.broadinstitute.org/morpheus) with using average linkage method and one minus Pearson correlation metric.

#### RNA sequencing of mouse pancreatic cancer cells

Murine PDAC cells were freshly isolated from dissociated pancreatic tumor tissues followed by FACS sorting (see Cell Lines). Total RNA was isolated using TRIzol (#15596-026, Life technologies) according to the manufacturer’s protocol. RNA from established cell lines was extracted with the use of an RNeasy Mini Kit (#74104, Qiagen). RNA samples were further submitted for RNAseq analysis. Total RNA libraries were prepared by using Pico Input SMARTer Stranded Total RNA-Seq Kit (Takara). In short, 250pg-10ng total RNA from each sample was reverse-transcribed via random priming and reverse transcriptase. Full-length cDNA was obtained with SMART (Switching Mechanism At 5’ end of RNA Template) technology. The template-switching reaction kept the strand orientation of the RNA. The ribosomal cDNA was hybridized to mammalian-specific R-Probes and then cleaved by ZapR. Libraries containing Illumina adapter with TruSeq HT indexes were subsequently pooled and loaded to the Hiseq 2500. Single end reads at 75bp were generated for gene expression analyses.

Sequencing reads were analyzed for quality issues using FastQC (S.Andrews, http://www.bioinformatics.babraham.ac.uk/projects/fastqc/). Reads were aligned to mouse genome (mm10) using TopHat2 (PMID: 23618408) and absolute gene counts were quantified using HTSeq (PMID: 25260700). The resulting gene counts were used as input for differential expression analysis between KPC and KPCN clones and primary cells using DESeq2 (PMID: 25516281). Genes that are differentially expressed were selected for subsequent downstream analysis for identification of biological pathways and ontologies using Gene Set Enrichment Analysis (GSEA) (p-value < 0.001) (PMID: 16199517) using MsigDB datasets (PMID: 26771021). To graphically represent the significantly enriched datasets (False Discovery Rate < 25%) Enrichment Map was used (PMID: 21085593).

#### Analysis of PDAC patient tumor tissues

The data of mRNA expression and patient outcomes data for human pancreatic ductal adenocarcinoma were extracted from publicly available data portals. RNA-Seq data for the TCGA PDAC tumor tissues were downloaded from Braod GDAC Firehose poratal (https://gdac.broadinstitute.org/) Subsequent analyses were limited to 76 cases with inferred PDAC fraction >30% by ESTIMATE method (Yoshihara et al., 2013) as described in (TCGA., 2017). The mRNA gene expression array data from 103 primary pancreatic ductal adenocarcinoma samples were obtained from GSE50827 and matched to survival information from the International Cancer Genome Consortium (ICGC) data portal (http://dcc/icgc.org). One case was excluded from the ICGC series due to early death. The PDAC cases in both, TCGA and ICGC series were classified as classic or basal according to a weighted gene expression algorithm (Moffitt et al., 2015). The RNA sequencing data from 85 human PDAC patient-derived xenografts was obtained from Champions Oncology, Inc., Hackensack, NJ, via TumorGraft® database (https://database.championsoncology.com/). The raw gene expression data used for analyzing subtype-specific pathway alterations in PDAC were normalized using Robust Multi-array Average (RMA) procedure (PMID: 12925520). Gene set variation analysis (GSVA, (Hanzelmann et al., 2013)) on the RMA normalized expression data was applied to identify pathways that are enriched in a single sample (method arguments: function=’gsva’; mx. Diff=TRUE; verbose=FALSE). Pathways that differ significantly between basal and classical cases were identified using Wilcoxon-Rank sum test using enrichment scores resulting from GSVA. Analyses to compare the overall-survival among the basal and classical subtypes were performed by comparing survival curves with log-rank tests. These were calculated using the R ‘survival’ package Therneau, T. M. & Grambsch, P. M. Modeling Survival Data: Extending the Cox Model (Springer-Verlag, 2010).

#### Wound healing assay

For the wound healing assay, cells were seeded in 12-well plates in concentration 2*10^5^ cells/ml/well in DMEM supplemented with 10% v/v FBS and 2mM L-glutamine with 100µg/ml Penicillin/Streptomycin and cultured to 95% confluence. The wound was created by scratching the cell monolayer with 200µl sterile pipette tip. To remove the debris and smooth the edge of the scratch, cells were washed once with the growth medium. Images of cells migrating into the wound were captured by time lapse microscopy (Nikon TE300 epi-fluorescent inverted microscope, 20x lens NA 0.45) every 15 minutes for duration of 20 hours at 37°C. Results were analyzed with ImageJ software.

#### TGFβ ELISA

The amount of TGFβ secreted by mouse pancreatic tumor cells was quantified by using the Mouse TGFβ 1 DuoSet ELISA (#DY1679-05, R&D systems), complemented with Sample Activation Kit 1 (#DY010, R&D systems) and DuoSet ELISA Ancillary Reagent Kit 1 (#DY007, R&D systems) according to the manufacturer’s instructions. Conditioned media for TGFβ secretion quantification was collected from KPC and KPCN tumor cells growing in DMEM supplemented with 1% v/v FBS and 2mM L-glutamine with 100µg/ml Penicillin/Streptomycin for 48 hours. Protein concentration of lysates, prepared from remaining cell pellets, was used for data normalization.

#### Confocal image acquisition

Confocal images were acquired with Leica TCS SP8 laser scanning confocal microscope with a 63x oil lens, NA 1.4. Images within each particular experiment were acquired under identical exposure conditions per channel.

#### Shh ELISA

The amount of Shh secreted by mouse pancreatic tumor cells was quantified by using the Mouse Shh-N ELISA Kit (#RAB0431, Millipore), according to the manufacturer’s instructions. KPC and KPCN cells were plated in DMEM supplemented with 10% v/v FBS and 2mM L-glutamine with 100µg/ml Penicillin/Streptomycin. The next day, when cell confluence achieved ∼90%, cells were washed twice with Hank’s Balanced Salt Solution (HBSS) and starved in HBSS overnight. Then fresh HBSS with/without Cholesterol Lipid Concentrate (#12531018, Gibco) was added and collected in 8 hours for analysis of the amount of secreted Shh. Protein concentration of lysates, prepared from remaining cell pellets, was used for data normalization.

#### mNSDHL expression in KPCN cells

In order to create KPCN cells expressing mouse NSDHL protein we used lentiviral construct pLEX-HA-MYC (#OHS4492, Thermo Scientific Open Biosystems). Lentivirus production was performed by co-transfecting HEK293T packaging cells with mNSDHL coding transfer vector and packaging plasmid mix included in Trans-Lentiviral ORF packaging kit with Calcium Phosphate (#TLP5916, Thermo Scientific Open Biosystems) according to the manufacturer’s protocol. KPCN cell lines were further infected with obtained lentiviruses and desired cells were selected by Puromycin (#P8833, Sigma) treatment (20µg/ml).

#### Cholesterol measurement

Cholesterol level in KPC and KPCN cells as well as in FBS and LDS containing media was measured by Amplex Red Kit (#A12216, Life Technologies), according to the manufacturer’s instructions.

#### siRNA transfection

siRNAs targeting *Tgfbr1*, *Acvr1b*, *Acvr1c* genes and control (*Gl2*) were obtained from Qiagen (#SI02735194, #SI01447040, #SI01447033, #SI01447019, #SI00888916, #SI00888909, #SI00888902, #SI00888895, #SI00888944, #SI00888937, #SI00888930, #SI00888923, #SI03650353, Qiagen). For control (*Gl2*) one siRNA was used and for targets of our interest (*Tgfbr1*, *Acvr1b*, *Acvr1c*) mix of four siRNAs per one gene was used. Cells were transfected with siRNA at 30 nM (total for four siRNAs) concentrations mixed with HiPerfect Transfection Reagent (#301704, Qiagen) according to the manufacturer’s reverse transfection protocol. In 24 hours after plating, cells were treated with media containing LDS or LDS with 1uM Compactin (#sc-200853, Santa Cruz). RNA and protein lysates were collected in 48 hours after treatment.

#### Phalloidin staining of stress fibers

To detect activation of stress fibers we performed phalloidin staining. KPC3 cells were plated on coverslips in DMEM supplemented with 10% v/v FBS and 2mM L-glutamine with 100µg/ml Penicillin/Streptomycin. The next day cells were washed with serum free medium and placed in DMEM supplemented with 2mM L-glutamine with 100µg/ml Penicillin/Streptomycin and 5% v/v FBS or 5% v/v LDS and 1uM compactin (#sc-200853, Santa Cruz). After 48 hours of incubation cells were starved for 4 hours in serum free medium. Then coverslips were fixed in 4%PFA and cells were permeabilized with 0.25% Triton X-100 (#BP151-100, Fisher Scientific), blocked in 3%BSA with 0.01% Triton X-100 and stained with Alexa Fluor 488 Phalloidin (#A12379, Invitrogen).

### STATISTICAL ANALYSIS

For analysis of continuous data we used Wilcoxon tests, and binary outcomes were compared using Fisher’s exact test. Repeated measures (i.e. multiple measures within mouse) were analyzed using generalized linear regression models with Generalized Estimating Equations (Liang and Zeger, 1986). Growth curves were modeled using linear regression with interactions between treatment and time, again using GEE to account for within-sample correlation. Survival time outcomes were assessed using Kaplan-Meier curves with log-rank tests.

### DATA AND SOFTWARE AVAILABILITY

RNAseq data reported in Figure 3 has been deposited into the Sequence Read Archive (SRA) database, deposition PRJNA530747 (https://www.ncbi.nlm.nih.gov/sra).

## References

Aiello, N. M., Brabletz, T., Kang, Y., Nieto, M. A., Weinberg, R. A., and Stanger, B. Z. (2017). Upholding a role for EMT in pancreatic cancer metastasis. Nature 547, E7–E8.

Aiello, N. M., Maddipati, R., Norgard, R. J., Balli, D., Li, J., Yuan, S., Yamazoe, T., Black, T., Sahmoud, A., Furth, E. E., et al. (2018). EMT Subtype Influences Epithelial Plasticity and Mode of Cell Migration. Developmental cell 45, 681–695 e684.

Aung, K. L., Fischer, S. E., Denroche, R. E., Jang, G. H., Dodd, A., Creighton, S., Southwood, B., Liang, S. B., Chadwick, D., Zhang, A., et al. (2018). Genomics-Driven Precision Medicine for Advanced Pancreatic Cancer: Early Results from the COMPASS Trial. Clinical cancer research : an official journal of the American Association for Cancer Research 24, 1344–1354.

Bailey, P., Chang, D. K., Nones, K., Johns, A. L., Patch, A. M., Gingras, M. C., Miller, D. K., Christ, A. N., Bruxner, T. J., Quinn, M. C., et al. (2016). Genomic analyses identify molecular subtypes of pancreatic cancer. Nature 531, 47–52.

Bang, U. C., Watanabe, T., and Bendtsen, F. (2018). The relationship between the use of statins and mortality, severity, and pancreatic cancer in Danish patients with chronic pancreatitis. Eur J Gastroenterol Hepatol 30, 346–351.

Bardeesy, N., Aguirre, A. J., Chu, G. C., Cheng, K. H., Lopez, L. V., Hezel, A. F., Feng, B., Brennan, C., Weissleder, R., Mahmood, U., et al. (2006). Both p16(Ink4a) and the p19(Arf)-p53 pathway constrain progression of pancreatic adenocarcinoma in the mouse. Proceedings of the National Academy of Sciences of the United States of America 103, 5947–5952.

Brown, M. S., Faust, J. R., Goldstein, J. L., Kaneko, I., and Endo, A. (1978). Induction of 3-hydroxy-3-methylglutaryl coenzyme A reductase activity in human fibroblasts incubated with compactin (ML-236B), a competitive inhibitor of the reductase. The Journal of biological chemistry 253, 1121–1128.

Chari, S. T., Kelly, K., Hollingsworth, M. A., Thayer, S. P., Ahlquist, D. A., Andersen, D. K., Batra, S. K., Brentnall, T. A., Canto, M., Cleeter, D. F., et al. (2015). Early detection of sporadic pancreatic cancer: summative review. Pancreas 44, 693–712.

Chen, M., Zhang, J., Sampieri, K., Clohessy, J. G., Mendez, L., Gonzalez-Billalabeitia, E., Liu, X. S., Lee, Y. R., Fung, J., Katon, J. M., et al. (2018). An aberrant SREBP-dependent lipogenic program promotes metastatic prostate cancer. Nature genetics 50, 206–218.

Chen, P. Y., Qin, L., Li, G., Tellides, G., and Simons, M. (2016). Smooth muscle FGF/TGFbeta cross talk regulates atherosclerosis progression. EMBO molecular medicine 8, 712–728.

Chen, W. C., Boursi, B., Mamtani, R., and Yang, Y. X. (2019). Total Serum Cholesterol and Pancreatic Cancer: A Nested Case-Control Study. Cancer epidemiology, biomarkers & prevention : a publication of the American Association for Cancer Research, cosponsored by the American Society of Preventive Oncology 28, 363–369.

Collisson, E. A., Sadanandam, A., Olson, P., Gibb, W. J., Truitt, M., Gu, S., Cooc, J., Weinkle, J., Kim, G. E., Jakkula, L., et al. (2011). Subtypes of pancreatic ductal adenocarcinoma and their differing responses to therapy. Nat Med 17, 500–503.

Cordenonsi, M., Dupont, S., Maretto, S., Insinga, A., Imbriano, C., and Piccolo, S. (2003). Links between tumor suppressors: p53 is required for TGF-beta gene responses by cooperating with Smads. Cell 113, 301–314.

Cunningham, D., DeBarber, A. E., Bir, N., Binkley, L., Merkens, L. S., Steiner, R. D., and Herman, G. E. (2015). Analysis of hedgehog signaling in cerebellar granule cell precursors in a conditional Nsdhl allele demonstrates an essential role for cholesterol in postnatal CNS development. Hum Mol Genet 24, 2808–2825.

David, C. J., Huang, Y. H., Chen, M., Su, J., Zou, Y., Bardeesy, N., Iacobuzio-Donahue, C. A., and Massague, J. (2016). TGF-beta Tumor Suppression through a Lethal EMT. Cell 164, 1015–1030.

Deng, Y. Z., Cai, Z., Shi, S., Jiang, H., Shang, Y. R., Ma, N., Wang, J. J., Guan, D. X., Chen, T. W., Rong, Y. F., et al. (2018). Cilia loss sensitizes cells to transformation by activating the mevalonate pathway. The Journal of experimental medicine 215, 177–195.

Di Guglielmo, G. M., Le Roy, C., Goodfellow, A. F., and Wrana, J. L. (2003). Distinct endocytic pathways regulate TGF-beta receptor signalling and turnover. Nat Cell Biol 5, 410–421.

Edlund, S., Landstrom, M., Heldin, C. H., and Aspenstrom, P. (2002). Transforming growth factor-beta-induced mobilization of actin cytoskeleton requires signaling by small GTPases Cdc42 and RhoA. Molecular biology of the cell 13, 902–914.

Fabregat, A., Sidiropoulos, K., Viteri, G., Forner, O., Marin-Garcia, P., Arnau, V., D’Eustachio, P., Stein, L., and Hermjakob, H. (2017). Reactome pathway analysis: a high-performance in-memory approach. BMC bioinformatics 18, 142.

Franco-Barraza, J., Francescone, R., Luong, T., Shah, N., Madhani, R., Cukierman, G., Dulaimi, E., Devarajan, K., Egleston, B. L., Nicolas, E., et al. (2017). Matrix-regulated integrin alphavbeta5 maintains alpha5beta1-dependent desmoplastic traits prognostic of neoplastic recurrence. Elife 6.

Freed-Pastor, W. A., Mizuno, H., Zhao, X., Langerod, A., Moon, S. H., Rodriguez-Barrueco, R., Barsotti, A., Chicas, A., Li, W., Polotskaia, A., et al. (2012). Mutant p53 disrupts mammary tissue architecture via the mevalonate pathway. Cell 148, 244–258.

Gabitova, L., Restifo, D., Gorin, A., Manocha, K., Handorf, E., Yang, D. H., Cai, K. Q., Klein-Szanto, A. J., Cunningham, D., Kratz, L. E., et al. (2015). Endogenous Sterol Metabolites Regulate Growth of EGFR/KRAS-Dependent Tumors via LXR. Cell reports 12, 1927–1938.

Genkinger, J. M., Kitahara, C. M., Bernstein, L., Berrington de Gonzalez, A., Brotzman, M., Elena, J. W., Giles, G. G., Hartge, P., Singh, P. N., Stolzenberg-Solomon, R. Z., et al. (2015). Central adiposity, obesity during early adulthood, and pancreatic cancer mortality in a pooled analysis of cohort studies. Ann Oncol 26, 2257–2266.

Golemis, E. A., Scheet, P., Beck, T. N., Scolnick, E. M., Hunter, D. J., Hawk, E., and Hopkins, N. (2018). Molecular mechanisms of the preventable causes of cancer in the United States. Genes & development 32, 868–902.

Guillaumond, F., Bidaut, G., Ouaissi, M., Servais, S., Gouirand, V., Olivares, O., Lac, S., Borge, L., Roques, J., Gayet, O., et al. (2015). Cholesterol uptake disruption, in association with chemotherapy, is a promising combined metabolic therapy for pancreatic adenocarcinoma. Proceedings of the National Academy of Sciences of the United States of America 112, 2473–2478.

Helliwell, S. B., Karkare, S., Bergdoll, M., Rahier, A., Leighton-Davis, J. R., Fioretto, C., Aust, T., Filipuzzi, I., Frederiksen, M., Gounarides, J., et al. (2015). FR171456 is a specific inhibitor of mammalian NSDHL and yeast Erg26p. Nature communications 6, 8613.

Hingorani, S. R., Wang, L., Multani, A. S., Combs, C., Deramaudt, T. B., Hruban, R. H., Rustgi, A. K., Chang, S., and Tuveson, D. A. (2005). Trp53R172H and KrasG12D cooperate to promote chromosomal instability and widely metastatic pancreatic ductal adenocarcinoma in mice. Cancer Cell 7, 469–483.

Hong, J. Y., Nam, E. M., Lee, J., Park, J. O., Lee, S. C., Song, S. Y., Choi, S. H., Heo, J. S., Park, S. H., Lim, H. Y., et al. (2014). Randomized double-blinded, placebo-controlled phase II trial of simvastatin and gemcitabine in advanced pancreatic cancer patients. Cancer chemotherapy and pharmacology 73, 125–130.

Hua, X., Wu, J., Goldstein, J. L., Brown, M. S., and Hobbs, H. H. (1995). Structure of the human gene encoding sterol regulatory element binding protein-1 (SREBF1) and localization of SREBF1 and SREBF2 to chromosomes 17p11.2 and 22q13. Genomics 25, 667–673.

Huang, B. Z., Chang, J. I., Li, E., Xiang, A. H., and Wu, B. U. (2017). Influence of Statins and Cholesterol on Mortality Among Patients With Pancreatic Cancer. Journal of the National Cancer Institute 109.

Jackson, E. L., Willis, N., Mercer, K., Bronson, R. T., Crowley, D., Montoya, R., Jacks, T., and Tuveson, D. A. (2001). Analysis of lung tumor initiation and progression using conditional expression of oncogenic K-ras. Genes & development 15, 3243–3248.

Kojima, Y., Acar, A., Eaton, E. N., Mellody, K. T., Scheel, C., Ben-Porath, I., Onder, T. T., Wang, Z. C., Richardson, A. L., Weinberg, R. A., and Orimo, A. (2010). Autocrine TGF-beta and stromal cell-derived factor-1 (SDF-1) signaling drives the evolution of tumor-promoting mammary stromal myofibroblasts. Proc Natl Acad Sci U S A 107, 20009–20014.

Kretzschmar, M., Doody, J., Timokhina, I., and Massague, J. (1999). A mechanism of repression of TGFbeta/ Smad signaling by oncogenic Ras. Genes & development 13, 804–816.

Liberzon, A., Birger, C., Thorvaldsdottir, H., Ghandi, M., Mesirov, J. P., and Tamayo, P. (2015). The Molecular Signatures Database (MSigDB) hallmark gene set collection. Cell systems 1, 417–425.

Miller, D. S. J., Bloxham, R. D., Jiang, M., Gori, I., Saunders, R. E., Das, D., Chakravarty, P., Howell, M., and Hill, C. S. (2018). The Dynamics of TGF-beta Signaling Are Dictated by Receptor Trafficking via the ESCRT Machinery. Cell reports 25, 1841–1855 e1845.

Moffitt, R. A., Marayati, R., Flate, E. L., Volmar, K. E., Loeza, S. G., Hoadley, K. A., Rashid, N. U., Williams, L. A., Eaton, S. C., Chung, A. H., et al. (2015). Virtual microdissection identifies distinct tumor- and stroma-specific subtypes of pancreatic ductal adenocarcinoma. Nature genetics 47, 1168–1178.

Morton, J. P., Timpson, P., Karim, S. A., Ridgway, R. A., Athineos, D., Doyle, B., Jamieson, N. B., Oien, K. A., Lowy, A. M., Brunton, V. G., et al. (2010). Mutant p53 drives metastasis and overcomes growth arrest/senescence in pancreatic cancer. Proceedings of the National Academy of Sciences of the United States of America 107, 246–251.

Nelson, E. R., Wardell, S. E., Jasper, J. S., Park, S., Suchindran, S., Howe, M. K., Carver, N. J., Pillai, R. V., Sullivan, P. M., Sondhi, V., et al. (2013). 27-Hydroxycholesterol links hypercholesterolemia and breast cancer pathophysiology. Science 342, 1094–1098.

Offield, M. F., Jetton, T. L., Labosky, P. A., Ray, M., Stein, R. W., Magnuson, M. A., Hogan, B. L., and Wright, C. V. (1996). PDX-1 is required for pancreatic outgrowth and differentiation of the rostral duodenum. Development 122, 983–995.

Ohlund, D., Handly-Santana, A., Biffi, G., Elyada, E., Almeida, A. S., Ponz-Sarvise, M., Corbo, V., Oni, T. E., Hearn, S. A., Lee, E. J., et al. (2017). Distinct populations of inflammatory fibroblasts and myofibroblasts in pancreatic cancer. The Journal of experimental medicine 214, 579–596.

Ozdemir, B. C., Pentcheva-Hoang, T., Carstens, J. L., Zheng, X., Wu, C. C., Simpson, T. R., Laklai, H., Sugimoto, H., Kahlert, C., Novitskiy, S. V., et al. (2014). Depletion of carcinoma-associated fibroblasts and fibrosis induces immunosuppression and accelerates pancreas cancer with reduced survival. Cancer Cell 25, 719–734.

Peterson, T. R., and Sabatini, D. M. (2005). eIF3: a connecTOR of S6K1 to the translation preinitiation complex. Molecular cell 20, 655–657.

Pitroda, S. P., Khodarev, N. N., Beckett, M. A., Kufe, D. W., and Weichselbaum, R. R. (2009). MUC1-induced alterations in a lipid metabolic gene network predict response of human breast cancers to tamoxifen treatment. Proceedings of the National Academy of Sciences of the United States of America 106, 5837–5841.

Rahib, L., Smith, B. D., Aizenberg, R., Rosenzweig, A. B., Fleshman, J. M., and Matrisian, L. M. (2014). Projecting cancer incidence and deaths to 2030: the unexpected burden of thyroid, liver, and pancreas cancers in the United States. Cancer research 74, 2913–2921.

Renner, I. G., Wisner, J. R., Jr., and Rinderknecht, H. (1983). Protective effects of exogenous secretin on ceruletide-induced acute pancreatitis in the rat. J Clin Invest 72, 1081–1092.

Rhim, A. D., Mirek, E. T., Aiello, N. M., Maitra, A., Bailey, J. M., McAllister, F., Reichert, M., Beatty, G. L., Rustgi, A. K., Vonderheide, R. H., et al. (2012). EMT and dissemination precede pancreatic tumor formation. Cell 148, 349–361.

Rhim, A. D., Oberstein, P. E., Thomas, D. H., Mirek, E. T., Palermo, C. F., Sastra, S. A., Dekleva, E. N., Saunders, T., Becerra, C. P., Tattersall, I. W., et al. (2014). Stromal elements act to restrain, rather than support, pancreatic ductal adenocarcinoma. Cancer Cell 25, 735–747.

Sabnis, A. J., and Bivona, T. G. (2019). Principles of Resistance to Targeted Cancer Therapy: Lessons from Basic and Translational Cancer Biology. Trends in molecular medicine 25, 185–197.

Scheel, C., Eaton, E. N., Li, S. H., Chaffer, C. L., Reinhardt, F., Kah, K. J., Bell, G., Guo, W., Rubin, J., Richardson, A. L., and Weinberg, R. A. (2011). Paracrine and autocrine signals induce and maintain mesenchymal and stem cell states in the breast. Cell 145, 926–940.

Silvente-Poirot, S., and Poirot, M. (2014). Cancer. Cholesterol and cancer, in the balance. Science 343, 1445–1446.

Stankic, M., Pavlovic, S., Chin, Y., Brogi, E., Padua, D., Norton, L., Massague, J., and Benezra, R. (2013). TGF-beta-Id1 signaling opposes Twist1 and promotes metastatic colonization via a mesenchymal-to-epithelial transition. Cell reports 5, 1228–1242.

Talebi, A., Dehairs, J., Rambow, F., Rogiers, A., Nittner, D., Derua, R., Vanderhoydonc, F., Duarte, J. A. G., Bosisio, F., Van den Eynde, K., et al. (2018). Sustained SREBP-1-dependent lipogenesis as a key mediator of resistance to BRAF-targeted therapy. Nature communications 9, 2500.

TCGA. (2017). Integrated Genomic Characterization of Pancreatic Ductal Adenocarcinoma. Cancer Cell 32, 185–203 e113.

Tuveson, D. A., Shaw, A. T., Willis, N. A., Silver, D. P., Jackson, E. L., Chang, S., Mercer, K. L., Grochow, R., Hock, H., Crowley, D., et al. (2004). Endogenous oncogenic K-ras(G12D) stimulates proliferation and widespread neoplastic and developmental defects. Cancer Cell 5, 375–387.

Wu, J., Jiao, Y., Dal Molin, M., Maitra, A., de Wilde, R. F., Wood, L. D., Eshleman, J. R., Goggins, M. G., Wolfgang, C. L., Canto, M. I., et al. (2011). Whole-exome sequencing of neoplastic cysts of the pancreas reveals recurrent mutations in components of ubiquitin-dependent pathways. Proceedings of the National Academy of Sciences of the United States of America 108, 21188–21193.

Ying, H., Kimmelman, A. C., Lyssiotis, C. A., Hua, S., Chu, G. C., Fletcher-Sananikone, E., Locasale, J. W., Son, J., Zhang, H., Coloff, J. L., et al. (2012). Oncogenic Kras maintains pancreatic tumors through regulation of anabolic glucose metabolism. Cell 149, 656–670.

Zhao, W., Prijic, S., Urban, B. C., Tisza, M. J., Zuo, Y., Li, L., Tan, Z., Chen, X., Mani, S. A., and Chang, J. T. (2016). Candidate Antimetastasis Drugs Suppress the Metastatic Capacity of Breast Cancer Cells by Reducing Membrane Fluidity. Cancer research 76, 2037–2049.

Zheng, X., Carstens, J. L., Kim, J., Scheible, M., Kaye, J., Sugimoto, H., Wu, C. C., LeBleu, V. S., and Kalluri, R. (2015). Epithelial-to-mesenchymal transition is dispensable for metastasis but induces chemoresistance in pancreatic cancer. Nature 527, 525–530.

## SUPPLEMENTARY REFERENCES

Beglyarova, N., Banina, E., Zhou, Y., Mukhamadeeva, R., Andrianov, G., Bobrov, E., Lysenko, E., Skobeleva, N., Gabitova, L., Restifo, D., et al. (2016). Screening of Conditionally Reprogrammed Patient-Derived Carcinoma Cells Identifies ERCC3-MYC Interactions as a Target in Pancreatic Cancer. Clin Cancer Res 22, 6153–6163.

Cunningham, D., DeBarber, A.E., Bir, N., Binkley, L., Merkens, L.S., Steiner, R.D., and Herman, G.E. (2015). Analysis of hedgehog signaling in cerebellar granule cell precursors in a conditional Nsdhl allele demonstrates an essential role for cholesterol in postnatal CNS development. Hum Mol Genet 24, 2808–2825.

Cunningham, D., Swartzlander, D., Liyanarachchi, S., Davuluri, R.V., and Herman, G.E. (2005). Changes in gene expression associated with loss of function of the NSDHL sterol dehydrogenase in mouse embryonic fibroblasts. J Lipid Res 46, 1150–1162.

Franco-Barraza, J., Francescone, R., Luong, T., Shah, N., Madhani, R., Cukierman, G., Dulaimi, E., Devarajan, K., Egleston, B.L., Nicolas, E., et al. (2017). Matrix-regulated integrin alphavbeta5 maintains alpha5beta1-dependent desmoplastic traits prognostic of neoplastic recurrence. Elife 6.

Hanzelmann, S., Castelo, R., and Guinney, J. (2013). GSVA: gene set variation analysis for microarray and RNA-seq data. BMC bioinformatics 14, 7.

Helliwell, S.B., Karkare, S., Bergdoll, M., Rahier, A., Leighton-Davis, J.R., Fioretto, C., Aust, T., Filipuzzi, I., Frederiksen, M., Gounarides, J., et al. (2015). FR171456 is a specific inhibitor of mammalian NSDHL and yeast Erg26p. Nat Commun 6, 8613.

Hruban, R.H., Adsay, N.V., Albores-Saavedra, J., Anver, M.R., Biankin, A.V., Boivin, G.P., Furth, E.E., Furukawa, T., Klein, A., Klimstra, D.S., et al. (2006). Pathology of genetically engineered mouse models of pancreatic exocrine cancer: consensus report and recommendations. Cancer Res 66, 95–106.

Kim, M.P., Evans, D.B., Wang, H., Abbruzzese, J.L., Fleming, J.B., and Gallick, G.E. (2009). Generation of orthotopic and heterotopic human pancreatic cancer xenografts in immunodeficient mice. Nat Protoc 4, 1670–1680.

Liang, K.Y., and Zeger, S.L. (1986). Longitudinal Data-Analysis Using Generalized Linear-Models. Biometrika 73, 13–22.

Moffitt, R.A., Marayati, R., Flate, E.L., Volmar, K.E., Loeza, S.G., Hoadley, K.A., Rashid, N.U., Williams, L.A., Eaton, S.C., Chung, A.H., et al. (2015). Virtual microdissection identifies distinct tumor- and stroma-specific subtypes of pancreatic ductal adenocarcinoma. Nat Genet 47, 1168–1178.

Morris, J.P.t., Cano, D.A., Sekine, S., Wang, S.C., and Hebrok, M. (2010). Beta-catenin blocks Kras-dependent reprogramming of acini into pancreatic cancer precursor lesions in mice. J Clin Invest 120, 508–520.

Schindelin, J., Arganda-Carreras, I., Frise, E., Kaynig, V., Longair, M., Pietzsch, T., Preibisch, S., Rueden, C., Saalfeld, S., Schmid, B., et al. (2012). Fiji: an open-source platform for biological-image analysis. Nat Methods 9, 676–682.

TCGA. (2017). Integrated Genomic Characterization of Pancreatic Ductal Adenocarcinoma. Cancer cell 32, 185–203 e113.

Yoshihara, K., Shahmoradgoli, M., Martinez, E., Vegesna, R., Kim, H., Torres-Garcia, W., Trevino, V., Shen, H., Laird, P.W., Levine, D.A., et al. (2013). Inferring tumour purity and stromal and immune cell admixture from expression data. Nature communications 4, 2612.

