## Supplementary figures and images for "Cholesterol deprivation induces TGFβ signaling to promote basal differentiation in pancreatic cancer" (from another version of this preprint: version 1, 2019-05-09)

### Supplemental Figure 1

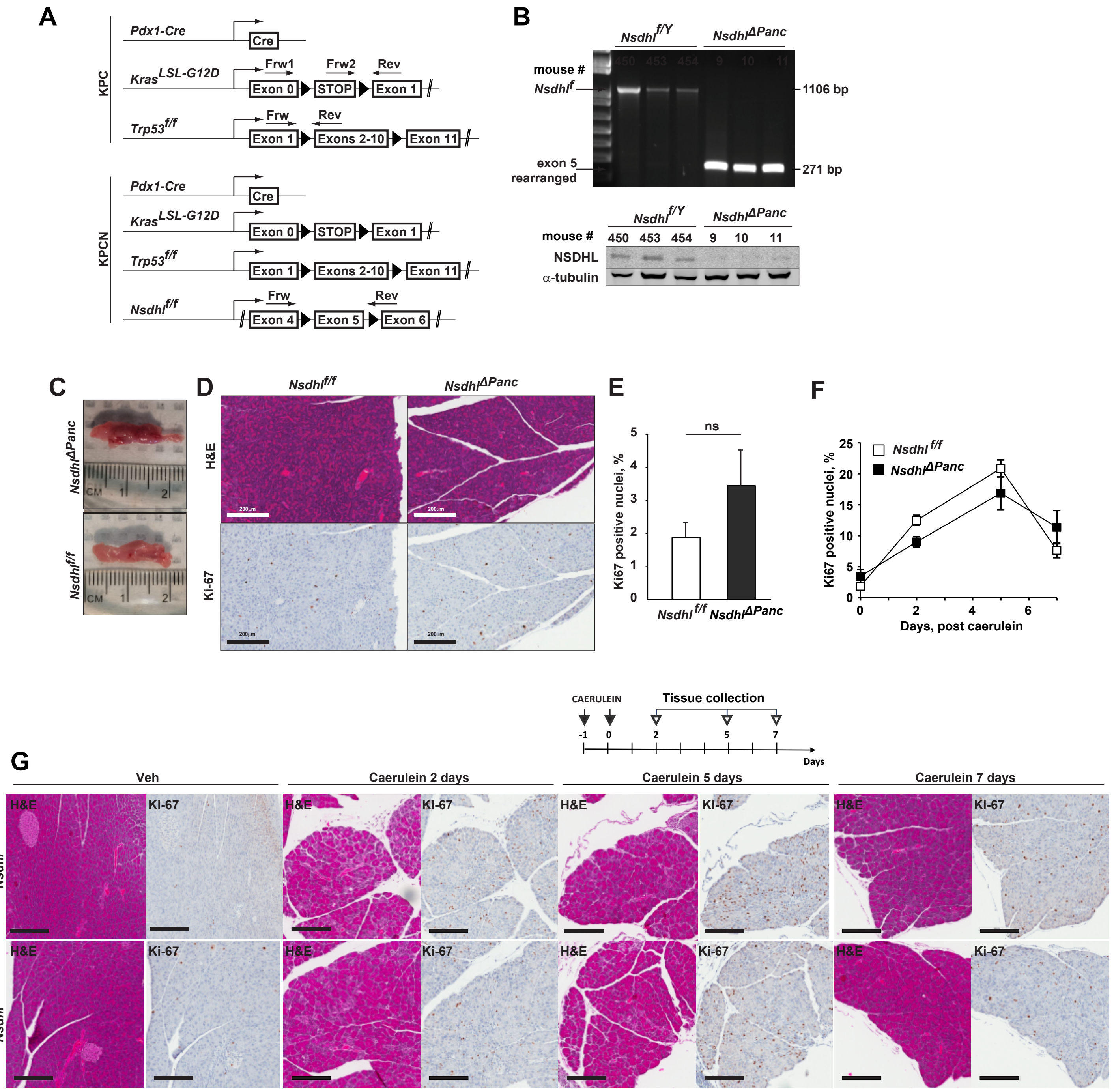

### Supplemental Figure 2

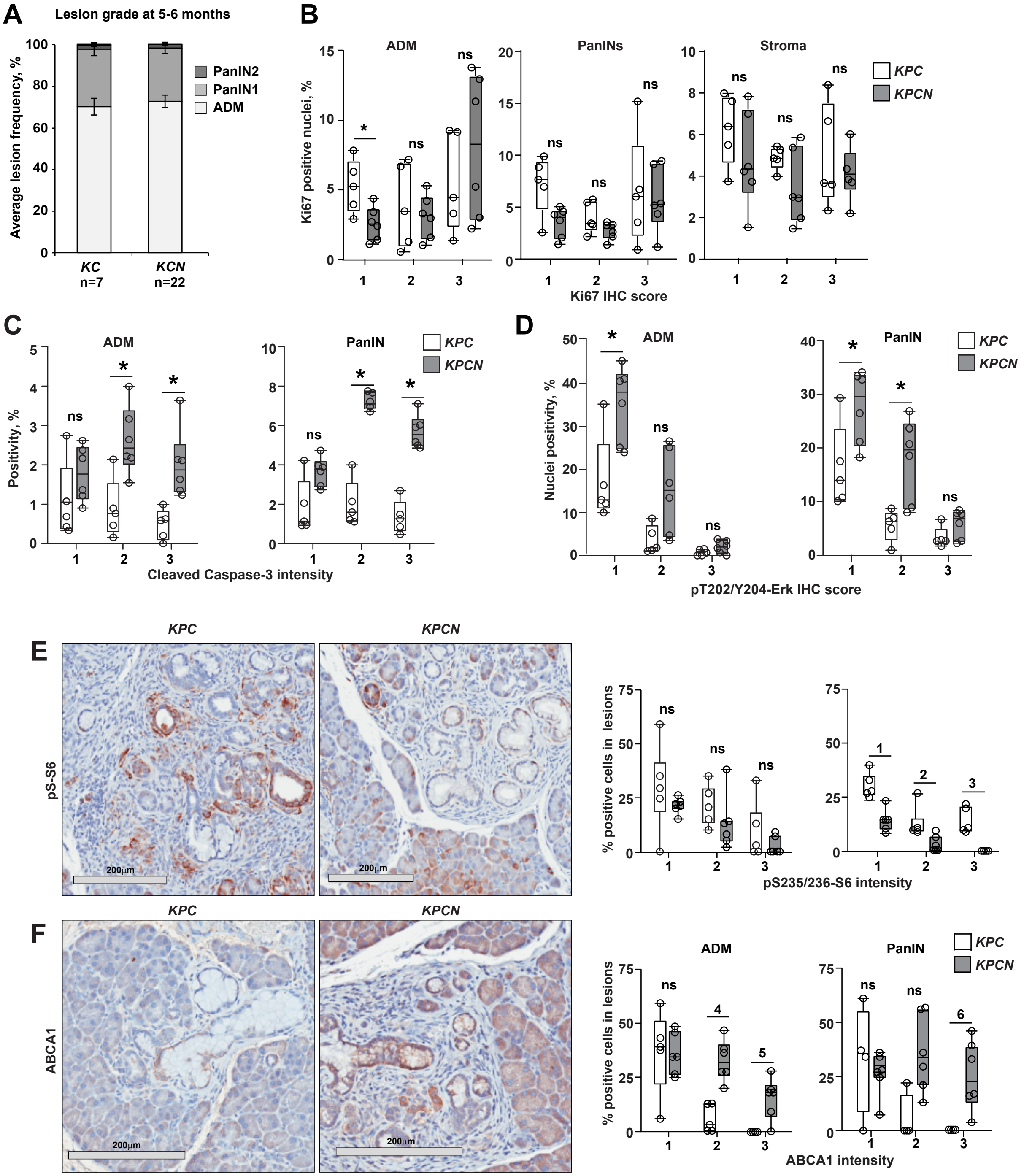

### Supplemental Figure 3

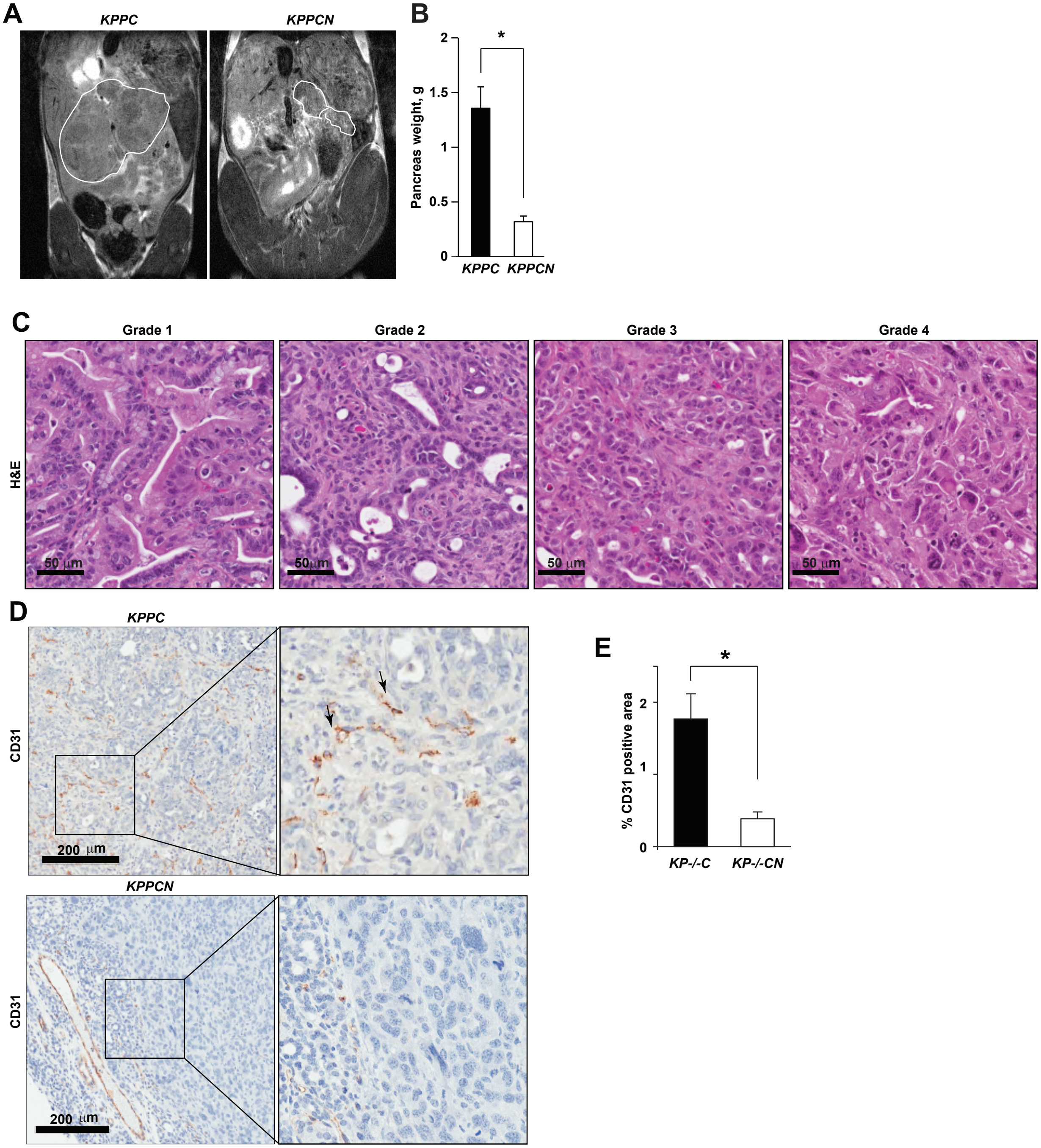

### Supplemental Figure 4

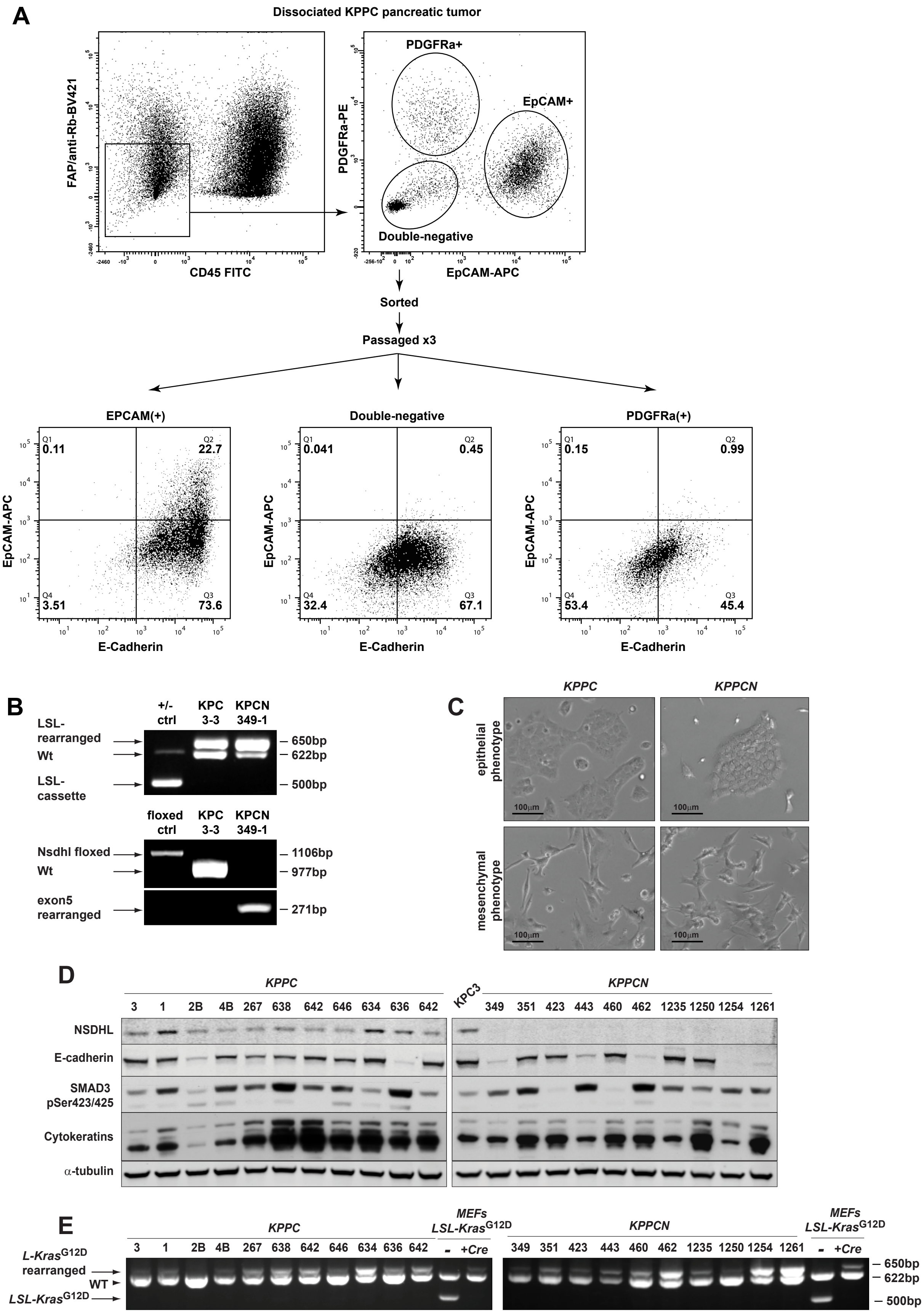

### Supplemental Figure 5

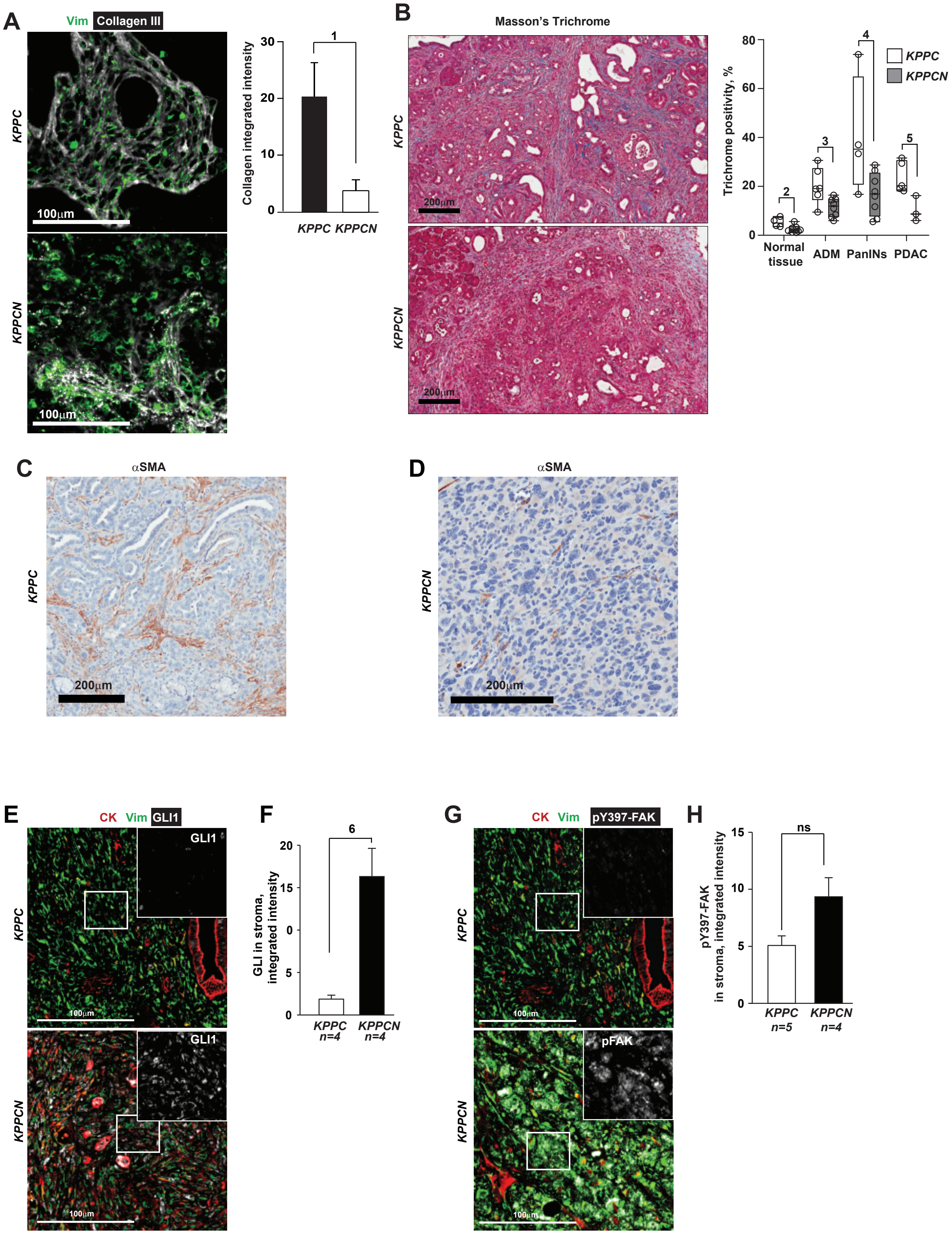

### Supplemental Figure 7

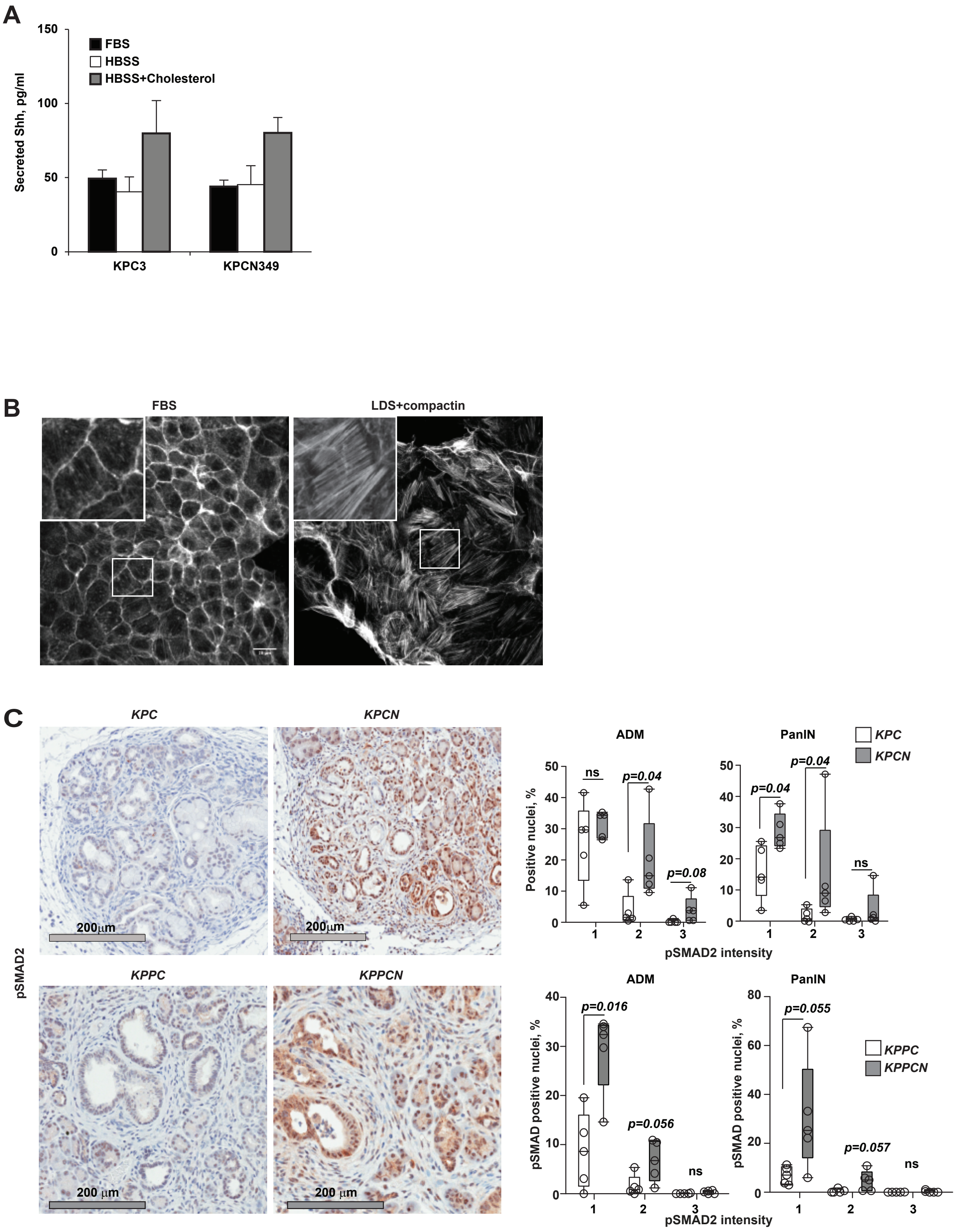
