## Supplemental Figure 6 for "Cholesterol deprivation induces TGFβ signaling to promote basal differentiation in pancreatic cancer" (from another version of this preprint: version 1, 2019-05-09)

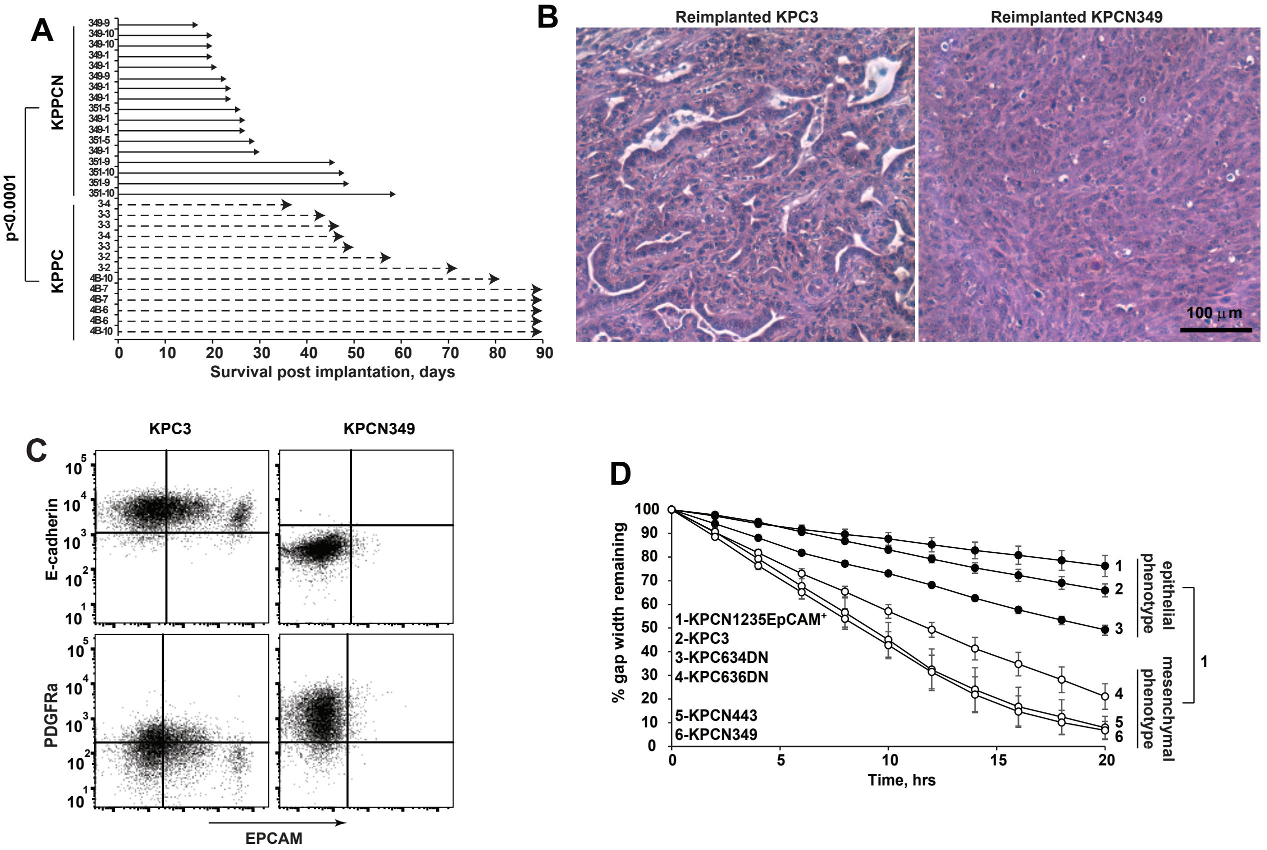


E

positive correlation, Pearson >0.45
negative correlation, Pearson <-0.45

| | CDH-1, WB | Cdh-1, qPCR | Zo-1, qPCR | Sox17, qPCR | Zeb2, qPCR | Grhl2, qPCR | Axin2, qPCR | Wnt7b, qPCR | Rnf43, qPCR | Wnt5a, qPCR | Wnt10b, qPCR | Tgfβ, qPCR | pSMAD3, WB | Tm7sf2, qPCR | Dhcr24, qPCR | Ebp, qPCR |
|---|---|---|---|---|---|---|---|---|---|---|---|---|---|---|---|---|
| CDH1, WB | 1 | | | | | | | | | | | | | | | |
| Cdh1 | 0.89 | 1 | | | | | | | | | | | | | | |
| Zo1 | 0.62 | 0.70 | 1 | | | | | | | | | | | | | |
| Sox17 | 0.52 | 0.47 | 0.57 | 1 | | | | | | | | | | | | |
| Zeb2 | -0.76 | -0.71 | -0.29 | -0.31 | 1 | | | | | | | | | | | |
| Grhl2 | 0.39 | 0.41 | 0.19 | 0.41 | -0.34 | 1 | | | | | | | | | | |
| Axin2 | -0.35 | -0.30 | 0.25 | -0.18 | 0.51 | -0.28 | 1 | | | | | | | | | |
| Wnt7b | 0.08 | 0.20 | 0.05 | -0.05 | -0.20 | 0.65 | -0.14 | 1 | | | | | | | | |
| Rnf43 | 0.59 | 0.59 | 0.56 | 0.25 | -0.47 | 0.54 | 0.04 | 0.30 | 1 | | | | | | | |
| Wnt5a | -0.46 | -0.45 | -0.25 | -0.18 | 0.84 | -0.20 | 0.28 | -0.23 | -0.37 | 1 | | | | | | |
| Wnt10b | -0.77 | -0.78 | -0.55 | -0.34 | 0.75 | -0.30 | 0.41 | -0.26 | -0.52 | 0.53 | 1 | | | | | |
| Tgfβ1 | -0.58 | -0.47 | 0.02 | -0.10 | 0.83 | -0.23 | 0.40 | -0.15 | -0.29 | 0.62 | 0.45 | 1 | | | | |
| pSMAD3, WB | -0.55 | -0.41 | -0.19 | -0.22 | 0.62 | -0.20 | 0.46 | 0.03 | -0.22 | 0.43 | 0.32 | 0.58 | 1 | | | |
| Tm7sf2 | 0.63 | 0.54 | 0.50 | 0.70 | -0.50 | 0.17 | -0.20 | -0.24 | 0.23 | -0.38 | -0.42 | -0.23 | -0.28 | 1 | | |
| Dhcr24 | 0.64 | 0.59 | 0.66 | 0.45 | -0.31 | 0.19 | 0.12 | -0.01 | 0.64 | -0.23 | -0.44 | -0.15 | 0.06 | 0.45 | 1 | |
| Ebp | 0.51 | 0.43 | 0.47 | 0.36 | -0.25 | -0.16 | -0.10 | -0.43 | 0.20 | -0.24 | -0.22 | 0.01 | -0.32 | 0.67 | 0.41 | 1 |

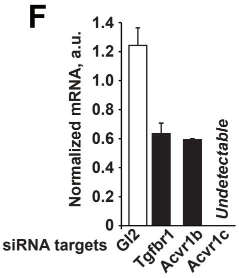
