## Supplementary material for "Cholesterol deprivation induces TGFβ signaling to promote basal differentiation in pancreatic cancer": Table S1 (from another version of this preprint: version 1, 2019-05-09)

**Supplementary Table S1.** Histological characterization of mouse adenocarcinoma.

| Sample# | PDAC histology | Age, days | Dominant PDAC component | PDAC grade | % area with solid growth |
| --- | --- | --- | --- | --- | --- |
|  | Genotype ***KPC***, n=24 |  |  |  |  |
| 13-5309 | Early adenocarcinoma | 129 | glandular | 1 | 0 |
| 13-5311 | Differentiated adenocarcinoma | 126 | glandular | 1 | 0 |
| 14-4602 | Differentiated adenocarcinoma | 142 | glandular | 1 | 0 |
| 14-4605 | Poorly differentiated adenocarcinoma | 197 | solid | 3 | 50 |
| 14-4612 | Differentiated adenocarcinoma | 154 | glandular | 1 | 10 |
| 14-4619 | Moderately differentiated adenocarcinoma | 122 | glandular | 2 | 0 |
| 14-4624 | Moderately differentiated adenocarcinoma | 153 | glandular | 2 | 20 |
| 15-0824 | Moderately differentiated adenocarcinoma | 170 | glandular | 2 | 10 |
| 15-0835 | Differentiated adenocarcinoma | 146 | glandular | 1 | 0 |
| 15-0847 | Moderately differentiated adenocarcinoma | 167 | glandular | 2 | 10 |
| 15-0848 | Cystic adenocarcinoma | 167 | glandular | 1 | 0 |
| 15-0859 | Differentiated adenocarcinoma | 101 | glandular | 1 | 0 |
| 15-0869 | Small focus, moderately differentiated adenocarcinoma | 90 | glandular | 2 | 10 |
| 15-0870 | Differentiated adenocarcinoma | 90 | glandular | 1 | 0 |
| 15-0871 | Moderately differentiated adenocarcinoma with necrosis | 136 | glandular | 2 | 10 |
| 16-1903 | Small focus of moderately differentiated adenocarcinoma | 141 | glandular | 2 | 0 |
| 16-1913 | Poorly differentiated adenocarcinoma with necrosis | 142 | glandular | 3 | 10 |
| 16-1917 | Poorly differentiated adenocarcinoma | 111 | solid | 3 | 50 |
| 16-1919 | Poorly differentiated adenocarcinoma | 111 | glandular | 3 | 50 |
| 16-1920 | Small focus of differentiated adenocarcinoma | 85 | glandular | 1 | 0 |
| 16-1925 | Differentiated adenocarcinoma | 112 | glandular | 1 | 0 |
| 16-1931 | Moderately differentiated adenocarcinoma | 111 | glandular | 2 | 10 |
| 16-1944 | Small focus, adenocarcinoma | 56 | glandular | 1 | 10 |
| 16-1958 | Moderately differentiated adenocarcinoma | 84 | glandular | 2 | 0 |
|  | Genotype ***KPCN***, n=3 |  |  |  |  |
| 17-0188 | Undifferentiated adenocarcinoma with sarcomatoid growth | 122 | solid | 4 | 70 |
| 17-5817 | Undifferentiated adenocarcinoma with sarcomatoid growth | 170 | solid | 4 | 90 |
| 18-5102 | Poorly differentiated adenocarcinoma | 191 | solid | 3 | <10% |
|  | Genotype ***KPPC***, n=15 |  |  |  |  |
| 15-2865 | Differentiated adenocarcinoma | 54 | glandular | 1 | 10 |
| 15-2866 | Differentiated adenocarcinoma | 51 | glandular | 1 | 0 |
| 15-2867 | Moderately differentiated adenocarcinoma | 51 | glandular | 2 | 10 |
| 16-3827 | Differentiated adenocarcinoma | 63 | glandular | 1 | 20 |
| 16-3829 | Moderately differentiated adenocarcinoma | 63 | glandular | 2 | 10 |
| 17-0022 | Differentiated adenocarcinoma | 32 | glandular | 1 | 10 |
| 17-0197 | Moderately differentiated adenocarcinoma | 66 | glandular | 2 | 10 |
| 17-0198 | Differentiated adenocarcinoma | 70 | glandular | 1 | <10 |
| 18-5014 | Differentiated adenocarcinoma | 64 | glandular | 1 | <10 |
| 18-5015 | Differentiated adenocarcinoma | 64 | glandular | 1 | 0 |
| 18-5019 | Poorly differentiated adenocarcinoma | 51 | glandular/solid | 3 | 30 |
| 18-5020 | Differentiated adenocarcinoma | 66 | glandular | 1 | 0 |
| 18-5098 | Differentiated adenocarcinoma | 54 | glandular | 1 | 0 |
| 18-5099 | Moderately differentiated adenocarcinoma | 61 | glandular | 2 | 30 |
| 18-5100 | Moderately differentiated adenocarcinoma | 65 | glandular | 2 | 20 |
|  | Genotype ***KPPCN***, n=25 |  |  |  |  |
| 16-3892 | Microfocus of undifferentiated adenocarcinoma | 56 | solid | 4 | 100 |
| 16-3910 | Microfocus of undifferentiated adenocarcinoma | 57 | solid | 4 | 100 |
| 16-3911 | Microfocus of undifferentiated adenocarcinoma | 57 | solid | 4 | 100 |
| 17-0020 | Undifferentiated adenocarcinoma | 82 | solid | 4 | 90 |
| 17-0036 | Undifferentiated adenocarcinoma | 94 | solid | 4 | 80 |
| 17-0179 | Undifferentiated adenocarcinoma | 86 | solid | 4 | 100 |
| 17-0183 | Microfocus of undifferentiated adenocarcinoma | 78 | solid | 4 | 100 |
| 17-0184 | Differentiated adenocarcinoma | 69 | glandular | 1 | 20 |
| 17-0185 | Undifferentiated adenocarcinoma with sarcomatoid growth | 84 | solid | 4 | 90 |
| 17-0186 | Undifferentiated adenocarcinoma | 75 | solid | 4 | 100 |
| 17-0189 | Undifferentiated adenocarcinoma | 80 | solid | 4 | 90 |
| 17-5809 | Poorly differentiated adenocarcinoma | 68 | glandular | 2 | 30 |
| 17-5814 | Undifferentiated adenocarcinoma with sarcomatoid growth | 78 | solid | 4 | 90 |
| 17-5815 | Undifferentiated adenocarcinoma with sarcomatoid growth | 83 | solid | 4 | 100 |
| 17-5822 | Differentiated adenocarcinoma | 86 | glandular | 1 | 20 |
| 17-5823 | Moderately differentiated adenocarcinoma | 71 | glandular | 2 | 30 |
| 17-6714 | Moderately differentiated adenocarcinoma | 57 | glandular | 2 | 10 |
| 18-5016 | Undifferentiated adenocarcinoma | 75 | solid | 4 | 70 |
| 18-5088 | Moderately differentiated adenocarcinoma | 86 | glandular | 2 | 30 |
| 18-5091 | Undifferentiated adenocarcinoma with sarcomatoid growth | 71 | solid | 4 | 100 |
| 18-5092 | Undifferentiated adenocarcinoma with sarcomatoid growth | 88 | solid | 4 | 100 |
| 18-5093 | Undifferentiated adenocarcinoma with sarcomatoid growth | 88 | solid | 4 | 100 |
| 18-5094 | Undifferentiated adenocarcinoma with sarcomatoid growth | 87 | solid | 4 | 90 |
| 18-5095 | Undifferentiated adenocarcinoma with sarcomatoid growth | 70 | solid | 4 | 80 |
| 18-5097 | Poorly differentiated adenocarcinoma | 77 | solid and glandular | 3&4 | 50 |

**Supplementary Table S2.** GSEA data murine PDAC. (excel file)

**Supplementary Table S3.** GSEA data PDAC PDX. (excel file)

**Supplementary Table S4.** GSEA data TCGA PDAC. (excel file)

**Supplementary Table S5.** Primers for genotyping and assays for qPCR.

| **Name** | **Sequence (5’ to 3’)** |
| --- | --- |
| Nsdhl exon5 rearrangement: Forward | gtg cta ctg tag act gaa cc |
| Nsdhl exon5 rearrangement: Reverse | gtg tcc ttg caa tct cag tg |
| Cre detection: oIMR1084 Forward | gcg gtc tgg cag taa aaa cta tc |
| Cre detection: oIMR1085 Reverse | gtg aaa cag cat tgc tgt cac tt |
| Internal control for Cre: oIMR7338 Forward | cta ggc cac aga att gaa aga tct |
| Internal control for Cre: oIMR7339 Reverse | gta ggt gga aat tct agc atc atc c |
| NSDHL flox5 detection: Forward | gtg cta ctg tag act gaa cc |
| NSDHL flox5 detection: Reverse | tct cct gga tgc tct gat ac |
| LSL-KRas detection and rearrangement: Forward 1 | gtc ttt ccc cag cac agt gc |
| LSL-KRas detection and rearrangement: Forward 2 | agc tag cca cca tgg ctt gag taa gtc tgc a |
| LSL-KRas detection and rearrangement: Reverse 1 | ctc ttg cct acg cca cca gct c |
| Trp53 flox detection: oIMR8543 Forward | ggt taa acc cag ctt gac ca |
| Trp53 flox detection: oIMR8544 Reverse | gga ggc aga gac agt tgg ag |
| **Name** |  |
| **SYBR Green Assays** | **Sequence (5’ to 3’)** |
| Grhl2 | For CCCATGACCTACCTCAACAAA  Rev CACCATCACCACACTCCTG |
| Wnt7b | For TCTCGTCGCTTTGTGGATG  Rev CGGTCCTCCAGAACCTTTC |
| Wnt10b | For GAATGCGGATCCACAACAAC  Rev CCCTCCAACAGGTCTTGAAT |
| Rnf43 | For AAGGGAAGCTAATGCAGTCC  Rev AGCTTGACGATGCTGATGAA |
| Sox17 | For TTTATGGTGTGGGCCAAAGA  Rev GCCTTCCAAGACTTGCCTAG |
| Axin2 | For GTCTCTACCTCATTTTCCGAGAA  Rev TCCAGCTCCAGTTTCAGTTT |
| Zo-1 | For ACCATGCCTAAAGCTGTCCCT  Rev TGTTAAAAATGCCACGAGCTGTAG |
| **Taqman Assays from Life Technologies** | **Cat#** |
| Zeb2 | Mm00497193_m1 |
| Cdh1 | Mm00486918_m1 |
| Tgfb1 | Mm03024053_m1 |
| Wnt5a | Mm00437347_m1 |
| Tm7sf2 |  |
| Dhcr24 |  |
| Ebp |  |
| Tgfbr2 |  |
