## Supplemental Data 1 for "Cholesterol deprivation induces TGFβ signaling to promote basal differentiation in pancreatic cancer" (from another version of this preprint: version 1, 2019-05-09)

**Supplementary figure legends.**

**Figure S1**. Development of *Nsdhl*-deficient murine pancreatic cancer model. (**A**) Schema of KPC and KPCN genetic constructs. *Thick arrowheads*, *loxP* sites. *Frw* and *Rev*, genotyping primers (see Supplementary Table S3). (**B**) PCR products and Western blot confirming rearrangement of *Nsdhl^flx^* genetic construct and depletion of NSDHL in pancreatic tissues of *Pdx1-Cre*-positive mice. (**C**) Normal size of *Nsdhl*-deficient pancreas.

(**D**) Hematoxylin-Eosin (H&E) and Ki67 staining of pancreatic tissues of *Nsdhl^f/f^* and *Nsdhl^∆Panc^* mice. Unperturbed basal proliferation (**E**), or reparative proliferation following acute caerulein-induced pancreatic injury (**F**) in pancreatic epithelial cells of *Nsdhl^f/f^* and *Nsdhl^∆Panc^* mice. (**G**) Representative H&E and Ki67 IHC images of pancreatic tissues of *Nsdhl^f/f^* and *Nsdhl^∆Panc^* mice following acute caerulein-induced pancreatic injury at indicated time points. Scale bars, 200 µm.

**Figure S2**. (**A**) Frequency per section of pre-neoplastic KRAS-transformed lesions in 5-6 months old *KC* and *KCN* mice. (**B-D**) Quantification of proliferating Ki67+ cells (**B**) in ADM and PanIN lesions and surrounding stroma, (**C**) cleaved caspase 3 and (**D**) phospho-ERK1/2 in *KPC* (n=5) and *KPCN* (n=6) mice at 5-6 months; *ns*, not significant, *, *p<*0.05. (**E**) IHC labeling of phosphorylated S235/236 ribosomal protein S6, and (**F**) ABCA1in age-matched 5-6-month old *KPC* (n=5) and *KPCN* (n=6) mice with quantification of IHC intensity in ADM, PanIN pancreatic epithelial lesions; scale bars, as indicated in µm. (1) *p*=0.004; (2) *p*=0.007; (3) *p*=0.004; (4) *p*=0.008; (5) *p*=0.02; (6) *p*=0.01, Wilcoxon test. In all figures, data are represented as mean±SEM.

**Figure S3.** (**A**) Micro-MRI images of littermate mice with outlined pancreatic tissues.

(**B**) Pancreas weights in *KPPC* and *KPPCN* mice at 7-8 weeks of age; *, *p*=0.0003.

(**C**) Representative Hematoxylin-Eosin images of different grades of PDAC. Scale bars, 60 µm. (**D**) Representative images of CD31 IHC staining of PDAC in *KPPC* and *KPPCN* mice. Scale bars, 200 µm. (**E**) Quantification of CD31 expression in *KPPC* and *KPPCN* pancreatic tumors. *, *p*<0.05.

**Figure S4.** Isolation and characterization of primary PDAC cells and PDAC cell lines.

(**A**) Representative plots of fluorescent activated cell sorting of single cell suspension obtained from dissociated *KPPC* tumor. (**B**) PCR products confirming rearrangement of *LSL-KRas^G12D^* and *Nsdhl^flx^* genetic constructs in indicated mouse PDAC cell lines. (**C**) Bright field images of epithelial and mesenchymal cells obtained as a result of sorting of *KPPC* and *KPPCN* tumors. Scale bars, 100 µm. (**D**) Western Blot of lysates obtained from *KPPC* and *KPPCN* clones illustrating expression of pancreatic adenocarcinoma differentiation markers. (**E**) Validation of PDAC cell lines for epithelial-specific genomic rearrangements of *LSL-Kras^G12D^* by PCR. DNA of NIH-3T3 fibroblasts was used as control.

**Figure S5**. Multicolor immunofluorescent characterization of pancreatic tumor tissues. (**A**) Deposition of dense collagen (white) in VIM+ (green) fibroblastic stroma is markedly reduced in *KPPCN* tumors. Scale bars, 100 µm. *Right*, quantification of collagen integrated intensity in *KPPCN* compared to *KPPC* adenocarcinoma. Data are represented as mean±SEM; (1), *p*=0.0043, Wilcoxon test. (**B**) Reduction of collagen deposition (blue) in *KPPCN* pancreatic tumors by Masson’s Trichrome staining. Scale bars, 200 µm. *Right*, quantification of percent of Trichrome positive (blue) area in *KPPCN* compared to *KPPC* adenocarcinoma. Data are represented as mean±SEM; (2), *p*=0.0283; (3), *p*=0.02; (4), *p*=0.0485; (5), *p*=0.0357, Wilcoxon test. (**C**) Representative image of alpha-smooth muscle actin (αSMA) staining of *KPPC* tumor tissue. (**D**) Representative image of alpha-smooth muscle actin (αSMA) staining of *KPPCN* tumor tissue. (**E**) Expression of GLI1 in fibroblastic stroma by multicolor immunofluorescence. (**F**) Quantification of GLI1 in restricted to CK-negative areas of the stroma in advanced *KPPCN* and *KPPC* pancreatic carcinomas; (3) *p*=0.0286, Wilcoxon test. (**G**) Expression of phospho-Y397-FAK (white) in fibroblastic stroma by multicolor immunofluorescence. Scale bars, 100 µm. (**H**) Quantification of pFAK in restricted to CK-negative areas of the stroma in advanced *KPPCN* (n=4) and *KPPC* (n=5) pancreatic carcinomas. Data represent mean±SEM of integrated intensity.

**Figure S6.** Functional characterization of murine PDAC cell lines. (**A**) Survival of mice with orthotopically re-implanted pancreatic tumors. Each arrow represents data from individual mouse that was injected with indicated cell line; *p*<0.0001; Wilcoxon test. Mice were followed for maximum of 90 days and euthanized in accord with the institutional guidelines. (**B**) Representative images of Hematoxylin-Eosin staining of pancreatic tumors obtained as a result of orthotopic implantation of *KPPC* (KPC3) and *KPPCN* (KPCN349) cell lines. Scale bars, 100 µm. (**C**) Expression of E-cadherin (CDH1), PDGFRα, and EpCAM in representative epithelial (*KPC3*) and mesenchymal (*KPCN349*) clones as determined by fluorescent activated cell sorting (FACS). (**D**) Clones with mesenchymal phenotype show increased motility in wound healing assay compare to epithelial clones; (1), *p*<0.0001, linear growth curve models on log-transformed volumes were modeled with generalized estimating equations. The p-value for the interaction between cell type and time tests for differences in the rate of change between cell types. (**E**) Pearson correlation coefficients for multiple parameters characterizing EMT of *KPPC* and *KPPCN* clones. (**F**) RT-PCR validation of siRNA silencing of TGFBR1, ACVR1B and ACVR1C in KPC3 cells.

**Figure S7.** Expanded data for Figure 4. (**A**) Sonic hedgehog measurements by ELISA in culture supernatants. Cells were grown to ~90% confluency, then serum starved in HBSS or given fresh 10% FBS-DMEM media for 18 hours. Cholesterol was added for 6 hours as indicated. All media was collected at the same time and subjected to ELISA. Graph is representative of 3 independent repeats. (**B**) qPCR validation of TGFBR2 expression in KPC3 cells under indicated culture conditions. (**C**) Patterns of F-actin stress fibers in KPC3 cells under indicated culture conditions as determined by Alexa 488-Phalloidin staining. *Scale bars*, 20 µM. (**D**) Activated TGFβ pathway signaling in NSDHL-deficient pancreatic lesions as assessed by phosphorylated SMAD2 immunohistochemistry. *Top*, 5-6 months old KPC and KPCN mice; *bottom,* 5 week old *KPPC* and *KPPCN* pancreatic lesions. *Right panel*, quantification of pSMAD2-positive nuclei in acinar-to-ductal metaplasia (ADM) and pancreatic intraepithelial neoplasm (PanIN) lesions represented as mean±SEM; *, *p*<0.05 by Wilcoxon test.
